# Imaging of plant calcium-sensor kinase conformation monitors real time calcium decoding *in planta*

**DOI:** 10.1101/2023.03.13.532409

**Authors:** Anja Liese, Bernadette Eichstädt, Sarah Lederer, Philipp Schulz, Jan Oehlschläger, José A Feijó, Waltraud X. Schulze, Kai R. Konrad, Tina Romeis

## Abstract

Changes in cytosolic calcium concentration are among the earliest reactions to a multitude of stress cues. Whereas a plethora of calcium-permeable channels may generate distinct calcium signatures and contribute to response specificities, the mechanisms by which calcium signatures are decoded is poorly understood. Here we develop a genetically encoded, FRET-based reporter that visualizes the conformational change of calcium-dependent protein kinases (CDPKs/CPKs), preceding kinase activation, for calcium-dependent AtCPK21 and calcium-independent AtCPK23. In pollen tubes, naturally displaying a physiological calcium range, CPK21-FRET, but not CPK23-FRET, report activity oscillations with similar features to cytosolic calcium, suggesting an isoform-specific calcium dependency and reversibility of the conformational change. In guard cells CPK21-FRET identifies CPK21 as a decoder of signal-specific calcium signatures in response to ABA and flg22. Based on this data, CDPK-FRET stands as a novel approach for tackling real-time live-cell calcium decoding in a multitude of plant developmental and stress responses.

## Introduction

Calcium (Ca^2+^) is the most important second messenger involved in signalling in practical all aspects of eukaryotic growth and development. The specificity of Ca^2+^ signalling is determined by the generation of Ca^2+^ concentration signals (encoding), followed by Ca^2+^ binding to proteins that convert Ca^2+^ signals into cellular responses (decoding) (Tian et al., 2020). Ca^2+^ signatures are defined through magnitude, number, duration and location of Ca^2+^ transients and are generated by the coordinated actions of Ca^2+^ channels and transporter (Yuan et al., 2014; Toyota et al., 2018; Tian et al., 2019; Thor et al., 2020; Tian et al., 2020; Bjornson et al., 2021; Köster et al., 2022; Xu et al., 2022). With respect to Ca^2+^ encoding major achievements in Ca^2+^ imaging have led to an increasing knowledge of real time Ca^2+^ dynamics in combination with Ca^2+^ permeable channel activities (Thor and Peiter, 2014; Toyota et al., 2018; Huang et al., 2019; Tian et al., 2019; Mou et al., 2020; Thor et al., 2020; Waadt et al., 2020; Bi et al., 2021; Bjornson et al., 2021; Eichstädt et al., 2021; Li et al., 2021; Guo et al., 2022; Tan et al., 2022; Xu et al., 2022). Yet, the Ca^2+^ decoding step, and in particular its assessment in real-time in living cells, remains a major challenge. Current effort focuses on *in situ* temporal and spatial activation of Ca^2+^ sensing proteins, with a view to directly record the Ca^2+^ signal information flow (decoding). CDPKs (CPKs in *Arabidopsis thaliana*) are Ca^2+^ sensor kinases specific to plants and important human parasites, binding Ca^2+^ directly and relaying the Ca^2+^ signal into protein phosphorylation (Harmon et al., 2000; Liese and Romeis, 2013; Simeunovic et al., 2016; Bender et al., 2018; Kudla et al., 2018; Yip Delormel and Boudsocq, 2019). In plants, the CDPK gene family is implicated in abiotic and biotic stress and in developmental signalling (Boudsocq et al., 2010; Geiger et al., 2010; Geiger et al., 2011; Gutermuth et al., 2013; Matschi et al., 2013; Brandt et al., 2015; Liu et al., 2017; Durian et al., 2020; Fu et al., 2022). CDPKs consist of a N-terminal variable domain, which may harbour myristoylation and palmitoylation membrane-localization motifs, followed by a serine/threonine protein kinase domain, a pseudosubstrate segment (PS), and a calmodulin-like domain (CLD) containing four consensus Ca^2+^ binding EF-hand motifs (Figure 1A) (Harmon et al., 2000; Liese and Romeis, 2013; Simeunovic et al., 2016; Bender et al., 2018; Kudla et al., 2018). CDPK activation can be considered as a two-step process: in the first step Ca^2+^ binding induces a conformational change which ‘opens’ the enzyme and which is prerequisite to the second step, the ATP-dependent phospho-transfer catalysed by the enzyme. In general, biochemical assessment of CDPK activities solely relies on step 2, evaluating enzyme activity by its catalytic trans-phosphorylation efficiency.

**Figure 1:**
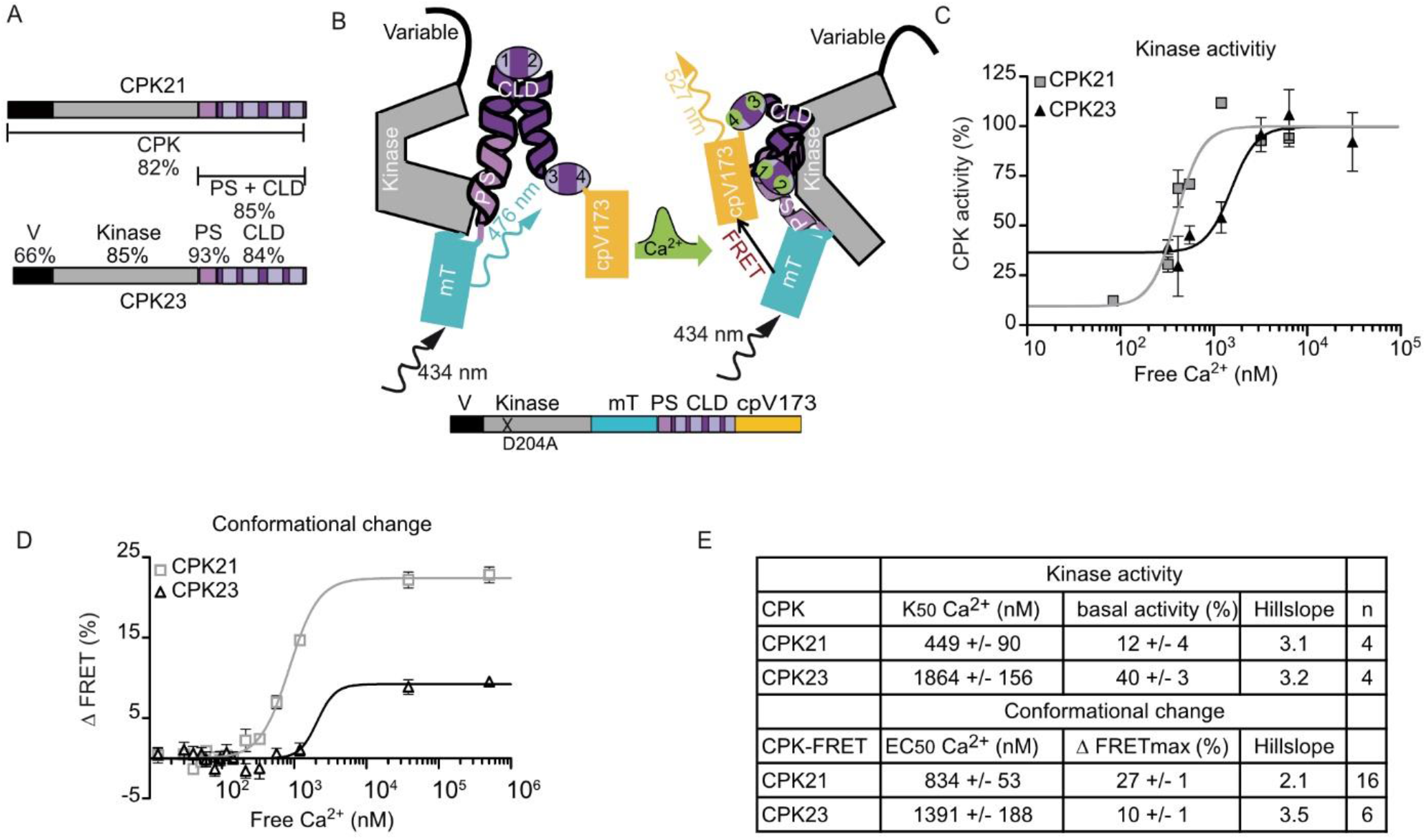
CDPK-FRET records CDPK isoform-specific Ca^2+^-dependencies of conformational change. A, Schemes of CPK21 and CPK23. Numbers indicate percentage of identical amino acids. A, B, V, variable domain; Kinase, kinase domain; PS, pseudosubstrate segment; CLD, calmodulin-like domain containing four EF-hand motifs (bright boxes). B, Scheme of CDPK-FRET between mTurquoise (mT) and cpVenus173 (cpV173) sandwiching PS-CLD and recording Ca^2+^-binding (green) induced conformational change. C, Kinase activity of CPK21 and CPK23 plotted against Ca^2+^-concentration (R^2CPK21^ = 0.91, R^2CPK23^ = 0.76). Activity is expressed as percentage of activity at full Ca^2+^-saturation (mean ± SEM n= 2-3, 7 Ca^2+^-concentration). D, FRET-recorded conformational change of CPK21 and CPK23 plotted against Ca^2+^-concentration (R^2CPK21^ = 0.98, R^2CPK23^ = 0.69). FRET efficiency change is given as percentage of increase in ratio over the baseline FRET signal (mean ± SEM n = 3-6, 15 Ca^2+^- concentrations). E, Summary of half-maximal kinase activity (K50) and half-maximal effective concentration (EC50) of conformational change in n independent experiments (mean ± SEM). The Hillslope is fitted as shared value for all data sets of the same enzyme.

We investigated two CDPKs from *Arabidopsis thaliana*, CPK21 and CPK23 which are implicated in plant abiotic stress signalling, where they participate in the ABA-mediated control of the stomatal aperture. Both enzymes are involved in the ABA-dependent activation of slow anion channel 1 (SLAC1) and SLAC1 homolog 3 (SLAH3) to mediate stomatal closure (Geiger et al., 2010; Geiger et al., 2011; Scherzer et al., 2012; Brandt et al., 2015). During this process, the Ca^2+^-dependent activation of SLAC1-type anion channels was predominantly assigned to CPK21 because CPK23 displayed a rather Ca^2+^ insensitive activity (Geiger et al., 2010; Scherzer et al., 2012). Remarkably, the same anion channels are also associated with stomatal closure in response to flg22, a pathogen-associated molecular pattern (PAMP) signal, as part of the plant pre-invasive immunity programme that prevents further pathogen invasion through open stoma (Guzel Deger et al., 2015; Wang and Gou, 2021).

The fact that flg22-induced stomatal closure on one hand involves Ca^2+^ signalling, and on the other hand SLAC1 type anion channels become activated as downstream recipients of Ca^2+^ signalling, pinpoints toward a role of CDPKs as prospective Ca^2+^ decoders (Thor and Peiter, 2014; Guzel Deger et al., 2015; Keinath et al., 2015; Thor et al., 2020).

Here, we developed a CDPK-associated ratiometric Förster resonance energy transfer (FRET) chimera that works as a sensor of the first step of the CDPK activation process, the Ca^2+^-dependent conformational change. Using the Ca^2+^-dependent *At*CPK21 and the Ca^2+^-independent *At*CPK23, we demonstrate that CDPK-FRET pairs genuinely record the conformational change in dependency of Ca^2+^, thus mirroring kinase activity. We calibrated these probes in the single cell model system of pollen tubes, and showed that the kinase activity closely mirrors the cytosolic Ca^2+^ oscillations during growth, supporting the notion of CDPK isoform-specific Ca^2+^-dependency as well as signal reversibility. Furthermore, we describe for the first time that CPK21 decodes both the ABA- and the flg22-induced Ca^2+^ signature in guard cells. Of relevance, the oscillatory features of Ca^2+^ kinetics and conformational change signal curves is suggestive of a biologically relevant distinction between sensing and decoding of Ca^2+^ signatures after elicitation with ABA or flg22.

## Results

### Novel FRET-based sensor records isoform-specific Ca^2+^-induced conformational change of CDPKs

The principle design for a CDPK conformation reporter, capable to record the enzyme’s Ca^2+^ binding-dependent conformational change, was deduced from *Toxoplasma gondii* CDPK1, whose X-ray structure indicates a significant Ca^2+^ binding-mediated conformational change for PS and CLD (Ojo et al., 2010; Wernimont et al., 2010). We therefore sandwiched PS-CLD between a FRET fluorescent protein pair, inserting mTurquoise (mT) as donor between kinase and PS domains, and Venus (circularly permutated at amino acid 173, cpV173) as acceptor C-terminal to the CLD (Figure 1B). If not stated otherwise kinase-deficient variants carrying an amino acid substitution in their ATP binding site were used for all CDPK-FRET fusion proteins to exclude an influence of auto-phosphorylation on the conformational change. The FRET-based CDPK conformation reporter was first established for *A. thaliana* CPK21, a Ca^2+^- sensitive CDPK with a half-maximal kinase activity (K50) of 449 nM Ca^2+^ (Geiger et al., 2010; Franz et al., 2011; Geiger et al., 2011) (Figure 1C, E). Different lengths and sequences of linkers connecting PS and CLD with the fluorophores plus a deletion variant lacking 8 C-terminal amino acids after the fourth EF hand were tested (Supplemental Figure S1A). All CPK21 conformation reporters, expressed and purified as recombinant fusion proteins, displayed enhanced FRET efficiency with increasing Ca^2+^ concentration and with similar half-maximal effective concentrations (EC50) for Ca^2+^ (Supplemental Figure S1A, B, F). The CPK21-FRET variant with the highest change in emission ratio (27 %, F4) was used as matrix for other CDPK based FRET constructs. The deduced EC50 value for the conformational change of CPK21-FRET is 834 nM Ca^2+^ (Figure 1D). To test if the mT insertion between kinase domain and PS influences kinase activity an additional CPK21_k_-FRET variant with an active kinase domain was generated. CPK21_k_-FRET displays low constitutive auto- and trans-phosphorylation activity and lacks a Ca^2+^-dependent increase in catalytic activity (Supplemental Figure S2). Thus, CDPK-FRET reports the conformational change in initial CDPK activation but may not monitor all regulatory aspects of biochemical CDPK catalytic activity that involves (auto/trans-) phosphorylation steps. Because EF-hands are known for a competitive binding of Mg^2+^ in addition to Ca^2+^ (Gifford et al., 2007), we assessed CPK21-FRET with Mg^2+^. At physiological free concentrations of 0.5 – 1 mM Mg^2+^ (Saris et al., 2000) decreased the CPK21-FRET efficiency only in the absence of Ca^2+^ (Supplemental Figure S1C), indicating binding selectivity for Ca^2+^ over Mg^2+^ and an different conformation when bound by Ca^2+^ or Mg^2+^. The Ca^2+^-induced changes of CPK21-FRET were stable over the plant cytosolic pH range of ∼ 7.2 – 7.5 (Felle, 2001; Zhou et al., 2021) (Supplemental Figure S1D). CPK23 displays a bi-partite pattern for catalytic kinase activity consisting of a Ca^2+^-independent core activity of 40 % and an additional Ca^2+^-dependent increase with low dependency (K50 = 1864 nM Ca^2+^) (Figure 1C). In accordance, CPK23-FRET reports a weak Ca^2+^-induced change in FRET efficiency of 10 % with a low Ca^2+^-dependency (EC50 = 1391 nM) (Figure 1D-E). Taken together these data demonstrate that CDPK-FRET conformational change measurements accurately report isoform-specific differences in Ca^2+^-dependency for CPK21 compared to CPK23.

### CDPK-FRET deciphers single amino acid residues as a determinant of Ca^2+^ dependent conformational change

CDPKs are characterized by their isoform-specific and highly variable Ca^2+^ dependencies for respective kinase activities (Geiger et al., 2010; Geiger et al., 2011; Boudsocq et al., 2012). To further assess the power and resolution of CDPK-FRET we employed the CDPK-FRET reporter to resolve the molecular basis for the contrasting Ca^2+^ dependencies of CPK21 and CPK23. The comparison of the primary sequences from these closely related enzymes (Figure 1A) identified an isoleucine at PS position 31 in CPK21 (I373) as conserved hydrophobic amino acid in the entire *A. thaliana* CDPK gene family, except in CPK23 where it is replaced by serine (S362) (Supplemental Fig S3A). Surrounding amino acids to S362 predict a CDPK phosphorylation motif (Huang et al., 2001). Indeed, upon transient expression in protoplasts, an (auto)-phosphorylated S362-containing peptide was detected in response to ABA when cells were transfected with active CPK23 but not with a kinase-deficient inactive CPK23 variant (Figure 2A, Supplemental Figure S3B). Mutual amino acid substitutions at PS position 31 were assessed for their impact on Ca^2+^- dependency of both, kinase activity and conformational change. In CPK21 substitutions at I373 for S (as in CPK23) and D (mimicking auto-phosphorylated CPK23) were investigated. In kinase assays these CPK21 substitutions at I373 to S or D caused an increase of the K50 value from 449 nM (WT) to 1111 nM (I373S) and 1905 nM (I373D) free Ca^2+^ (Figure 2B). In corresponding CPK21-FRET conformational change measurements an increase of the EC50 value from 834 nM (WT) to 925 nM (I373S) and 1701 nM (I373D) free Ca^2+^ was observed (Figure 2C). These data indicate that a single amino acid exchange is sufficient to turn the highly Ca^2+^ dependent CPK21 into a rather Ca^2+^ independent CDPK, resembling CPK23. Also, these data verify that CDPK-FRET reports Ca^2+^ dependencies with a similar resolution as catalytic activities measurements. *Vice versa*, in CPK23, the substitution S362I, renders the enzyme more sensitive toward Ca^2+^ in kinase assays, while S362D leads to constitutive Ca^2+^ independent activity (Supplemental Figure S3E). Interestingly, when employing CPK23-FRET, both introduced amino acid substitutions displayed a similar weak conformational change, with Ca^2+^ dependencies comparable to the native CPK23 protein (Supplemental Figure S3C). In these specific experiments the kinase-active form of CPK23_k_-FRET was applied to enable potential auto-phosphorylation at S362, whereby the active and inactive CPK23 FRET variants displayed similar half-maximal effective concentrations (EC50) and FRET efficiencies (Supplemental Figure S3D). To exclude that the comparatively low FRET efficiency of CPK23 may masks subtle effects, a CLD domain swap from CPK21 to CPK23 was conducted (Figure 2D). Chimeric CPK23CLD21–FRET shows increased maximal FRET efficiency (Figure 2F), indicating altered (higher) distance changes between donor and acceptor in dependency of CDPK conformation. Whereas the (low) Ca^2+^-dependency of CPK23-CLD21-FRET is comparable to native CPK23-FRET, remarkably, CPK23S362I-CLD21–FRET became Ca^2+^ dependent and displayed a Ca^2+^-dependent conformational change indistinguishable from CPK21 (Figure 2F). Also, when looking at kinase activities CPK23CLD21 and more pronounced CPK23S362I-CLD21 developed higher Ca^2+^ dependencies compared to independent CPK23 (Figure 2E). Taken together, these data uncover a single CPK23 auto-phosphorylation site as key to modify Ca^2+^-dependency and shift catalytic activity. In addition, CDPK-FRET has been established and validated as a novel tool to assess CDPK biochemical properties and activation independently, by using the first step of catalysis, the Ca^2+^ binding-dependent conformational change, as molecular readout.

**Figure 2:**
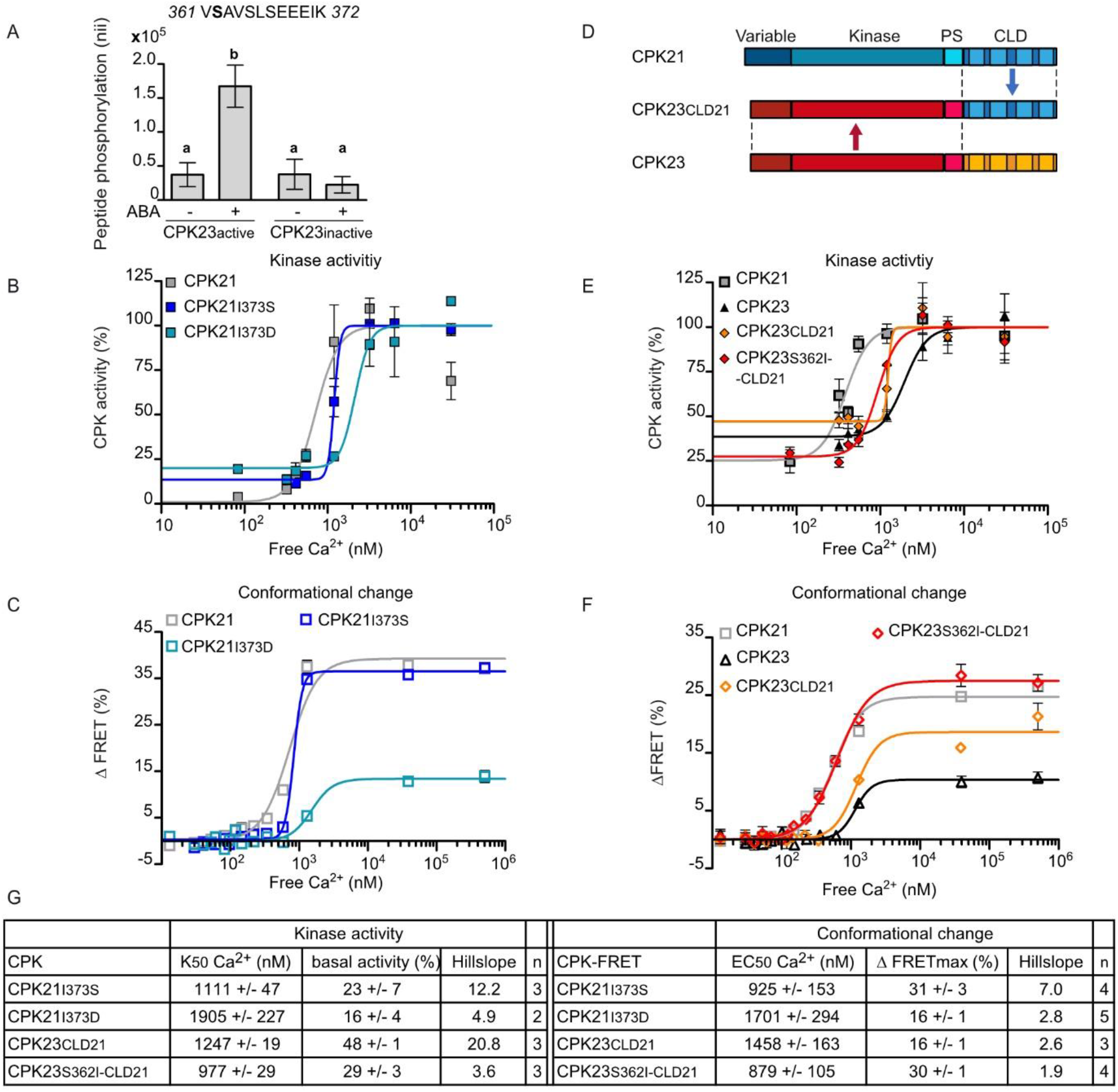
Unique single auto-phosphorylation site in pseudosubstrate segment controls Ca^2+^- dependency of CDPK conformational change and kinase activity. A, Arabidopsis protoplasts expressing catalytic active CPK23 or inactive CPK23 were treated with ABA, and *in vivo* phosphorylation at S362 was quantified via SRM mass spectrometry. Mean ± SEM combining three experiments p ≤ 0.05, one-way ANOVA, Tukey’s post-test, different letters indicate significant differences. B, Kinase activity of CPK21 and PS variants carrying an amino acid substitution (CPK21I373S, CPK21I373D) plotted against Ca^2+^-concentration (R^2CPK21^ = 0.85, R^2CPK21I373S^ = 0.97, R^2CPK21I373D^ = 0.87). C, FRET-recorded conformational change of CPK21, CPK21I373S, and CPK21I373D plotted against Ca^2+^-concentrations (R^2CPK21^ = 0.96, R^2CPK21373S^ = 0.99, R^2CPK21I373D^ = 0.88). D, Scheme of CPK23CLD21 chimeras. Abbreviation as in Figure 1A. E, F, Kinase activity (E) and FRET efficiency (F) of CPK21, CPK23, CPK23CLD21 and CPK23S362I-CLD21 plotted against Ca^2+^-concentrations (Kinase activity: R^2CPK21^ = 0.85, R^2CPK23^ = 0.89, R^2CPK23CLD21^ = 0.69, R^2CPK23S362I-CLD21^ = 0.92; Conformational change: R^2CPK21^ = 0.99, R^2CPK23^ = 0.81, R^2CPK23-CLD21^ = 0.91, R^2CPK23S362I-CLD21^ = 0.98). Kinase activity (B, E mean ± SEM n=2-4, 7-8 Ca^2+^-concentrations) and FRET-imaged conformational change (C, F mean ± SEM n= 3-6, 14-15 Ca^2+^-concentrations) are determined as described in Figure 1C and 1D. G, Summary of half-maximal kinase activity (K50) and half-maximal effective concentration (EC50) of conformational change in n independent experiments (mean ± SEM). The Hillslope is fitted as shared value for all data sets of the same enzyme.

### CDPK-FRET mirrors the cytosolic Ca^2+^ oscillatory pattern during pollen tube growth

Next, we aimed for the application of CDPK-FRET in an environment of changing Ca^2+^ concentrations in a plant cell system. Pollen tubes are an optimal single cell model system for cytosolic Ca^2+^ imaging, characterized by a tip-focused standing gradient frequently showing well defined oscillations during pollen tube growth (Michard et al., 2008; Konrad et al., 2011; Damineli et al., 2017; Michard et al., 2017; Li et al., 2021). Cytosolic CDPK-FRET variants were generated by ablating the myristoylation G2A and palmitoylation C3V motifs. These were transiently expressed in tobacco pollen that express as stable transgene the Ca^2+^ sensor R-GECO1 (red fluorescent genetically encoded Ca^2+^ indicator for optical imaging) (Zhao et al., 2011). CDPK-FRET and R-GECO1 signals were recorded simultaneously by time-lapse imaging of the transfected pollen. Of relevance, cytosolic Ca^2+^ concentrations in pollen tubes of tobacco has been quantified (Michard et al., 2008), and found to change from 0.2 to > 1.0 µM from the shank to the tip. Furthermore, during oscillations, values at the clear zone bellow the tip were determined to vary from 0.5 to > 1.0 µM. These values allow to estimate the Ca^2+^ changes captured by CPK21 and 23 FRET pairs with their distinct Ca^2+^- dependencies (Figure 1E). Accordingly, CPK21-FRET pollen displayed FRET intensity changes, which were in amplitude, phase and shape, closely mirroring the tip-focused cytosolic Ca^2+^ patterns, indicating a rapid reversibility of CPK21 conformational change (Figure 3A and Supplemental Figure S4, Supplemental movie S1). A high correlation coefficient (0.7 ± 0.2, Figure 3C-D) was obtained in synchronization analyses between the R-GECO1 monitored cytosolic Ca^2+^ concentration and the CDPK-FRET monitored CPK21 conformational change. In contrast, CPK23-FRET was unable to monitor any tip-focused oscillatory cytosolic Ca^2+^ concentration pattern, consistent with a low Ca^2+^- dependency of CPK23 kinase activity and small conformational changes (Figure 3B, Supplemental Figure S5 Supplemental movie S2). This is also evident from the weak correlation between Ca^2+^ changes and CPK23-FRET signal changes (correlation coefficient of 0.3 ± 0.1) (Figure 3C-D). In conclusion, cytosolic CPK21-FRET captures the oscillatory Ca^2+^ concentration pattern in pollen with a high correlation, in phase, and with no time delay (Figure 3D-E), thus demonstrating that it is a fast and accurate reporter/ sensor of Ca^2+^ changes.

**Figure 3:**
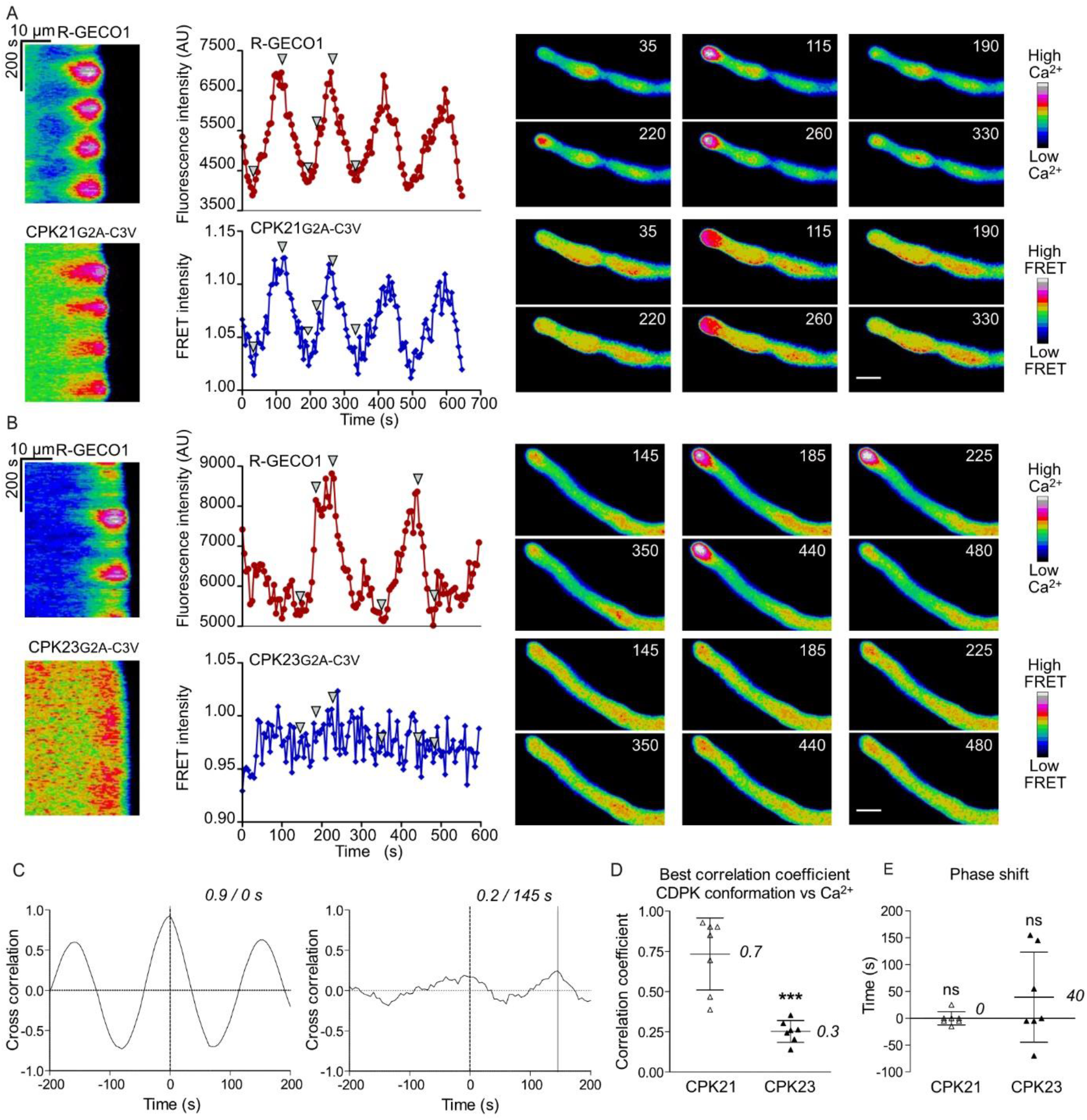
Tobacco pollen tube tip-focussed Ca^2+^-oscillations are decoded in real-time by CPK21-FRET but not by CPK23-FRET. A, B, Tobacco pollen carrying the R-GECO1 Ca^2+^-sensor as a stable transgene were transiently transformed with CDPK-FRET reporters CPK21G2A-C3V (A) or CPK23G2A-C3V (B). Growing pollen tubes were imaged in parallel for cytosolic changes in Ca^2+^-concentrations (top graphs, red) and CDPK conformational change (lower graphs, blue). Images were taken every 5 s for 645 s (A) and 660 s (B). Results are presented both as kymographs and as intensity-over-time-plots. Arrowheads in the intensity plots correspond to the false colored pollen tube images shown on the right side. Time in the pollen tube images is indicated in seconds and scale bar represents 15 μm. C, Synchronisation analysis between cytosolic Ca^2+^-concentration and CDPK conformational change in pollen tubes. The correlation coefficient between the R-GECO1 and CDPK-FRET signals is plotted for time delays of -200 s to + 200 s (CPK21 left panel, corresponding to 3A; CPK23, right panel, corresponding to 3B). A solid grey line marks the time delay with the highest correlation coefficient, which is indicated above the graph. The phase relationship between the signals is indicated as time difference in s. Time < 0 corresponds to leading, time > 0 to lagging R-GECO1 (Ca^2+^-concentration) signals, and 0 indicates no delay. Summary of synchronization analysis correlation coefficients (D) or time delay for the most probable match (in seconds) (E) are shown. Data represents the mean ± SD (numbers represent means) and dots represent the individual measurements (n = 7). D, A T-test yields a significant difference *** p ≤ 0.001 between correlation coefficients of either CPK21 or CPK23 conformational change to Ca^2+^- concentration. E, One-sided t-test reveals no significant differences (ns, p ≤ 0.05) to zero.

### CPK21-FRET identifies functional stress-specific Ca^2+^ decoding in guard cells

Guard cells respond to external application of either ABA or flg22 with cytosolic Ca^2+^ changes, which are relayed via different Ca^2+^ sensor kinases on anion channels as mediators of stomatal closure (Geiger et al., 2010; Geiger et al., 2011; Thor and Peiter, 2014; Brandt et al., 2015; Guzel Deger et al., 2015; Keinath et al., 2015). To test if CPK21 is biochemically activated in response to one or to both stimuli, stable *Arabidopsis* lines were generated in the Col-0 R-GECO1 background (Waadt et al., 2017) expressing plasma membrane localised CPK21-FRET under the control of a ß- estradiol-inducible promoter. After ABA or flg22 application to epidermal peels CPK21-FRET and R-GECO1 signals were recorded simultaneously by time-lapse imaging of selected single guard cells. Both stimuli induced an increase of cytosolic Ca^2+^- concentrations in the form of repetitive transients in the R-GECO1 signal, indicative for distinct Ca^2+^ signatures with a higher maximum signal change observed upon flg22 than ABA (Figure 4, Figure 5A). Remarkably, both stimuli also trigger a CPK21 conformational change as recorded by CPK21-FRET. Thus, CPK21 decodes and is activated not only by ABA, as one may have presumed (Geiger et al., 2010; Geiger et al., 2011), but also by flg22 (Figure 4 Supplemental Figure S6, S7, S8). Synchronization analysis shows that, for both stimuli Ca^2+^ concentration changes and respective CPK21 conformational changes are linked (correlation coefficient: ABA = 0.6 ± 0.1, flg22 = 0.7 ± 0.1, Figure 4, 5B). However, detailed evaluation of the Ca^2+^ signatures (curves shown in red) in relation to the FRET-monitored CPK21 conformational change signal curves (shown in blue) uncover stimulus-specific differences in the decoding of Ca^2+^ by CPK21. Of relevance, and in contrast to the analysis in pollen, where CPK21-FRET and R-GECO1 recorded almost identical kinetics and spatial pattern (∆area 6 ± 7 %, n = 3; Supplemental Figure S9), in the biological context of guard cells differences between Ca^2+^ concentration change and CPK21 conformational change kinetics are clearly visible (Figure 5C, blue shaded area in Figure 5E-F). This indicates that Ca^2+^ decoding is not synonymous to Ca^2+^ sensing. Interestingly, quantification of the difference in the ‘area under the curve’ between the plots of Ca^2+^ concentration changes and CPK21-FRET signal changes (blue shaded area in Figure 5E-F) yields a statistically significant higher ∆area for flg22 (46 ± 10 %) than for ABA (34 ± 10 %) (Figure 5D).

**Figure 4:**
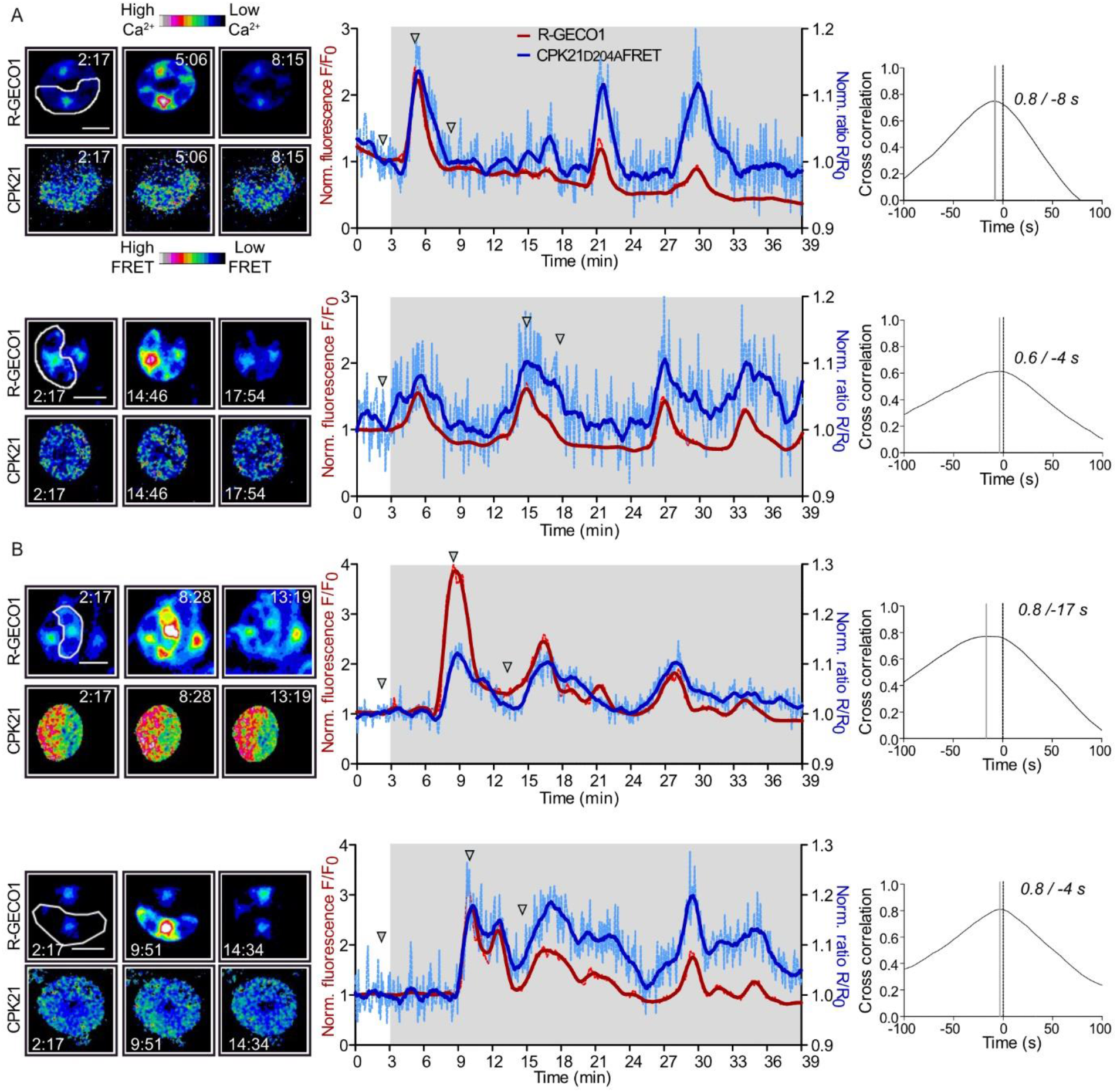
In guard cells CPK21-FRET uncovers decoding of ABA- and flg22-dependent Ca^2+^-signature *in vivo* in real-time. Parallel imaging of changes in the cytosolic Ca^2+^ concentration (red curve) and FRET ratio changes (blue curve) in response to 20 µM ABA (A) or 100 nM flg22 (B). Fluorescence images of measured guard cells (left panels) and intensity-over-time plots (middle panels) are shown. Continuous lines represent smoothed fluorescence intensity data (averaging 15 values on each side using a second order polynomial) and dotted lighter coloured lines represent normalized, original data. Images were taken every 4.16 s. The underlying grey area indicates the time interval of recording after ABA or flg22 treatment at 3 min. Micrographs represent selected time points indicated by arrowheads in the intensity plots. Time stamps are in the format mm:ss and scale bar represents 10 μm. The regions of interest (ROIs) used to measure signal intensity changes are framed white (leftmost panel). Quantification of phase relationships via cross correlation from ABA treated (A) and flg22 treated (B) guard cells are shown in the right-hand panels. Synchronisation analysis are based on adjusted signal changes (adjusted for signal decreases derived from technical artefacts) via dividing normalized signals by the trend line. The artificial trend line was calculated as linear regression of all data points. The correlation coefficient between the R-GECO1 and CDPK-FRET signals is plotted for time delays of -100 s to + 100 s. A solid grey line marks the time delay with the highest correlation coefficient. Time differences between the solid grey and dotted vertical (0 s) lines corresponds to leading (shift to left side) of R-GECO1 signals. Time delay and highest correlation coefficient are indicated.

**Figure 5:**
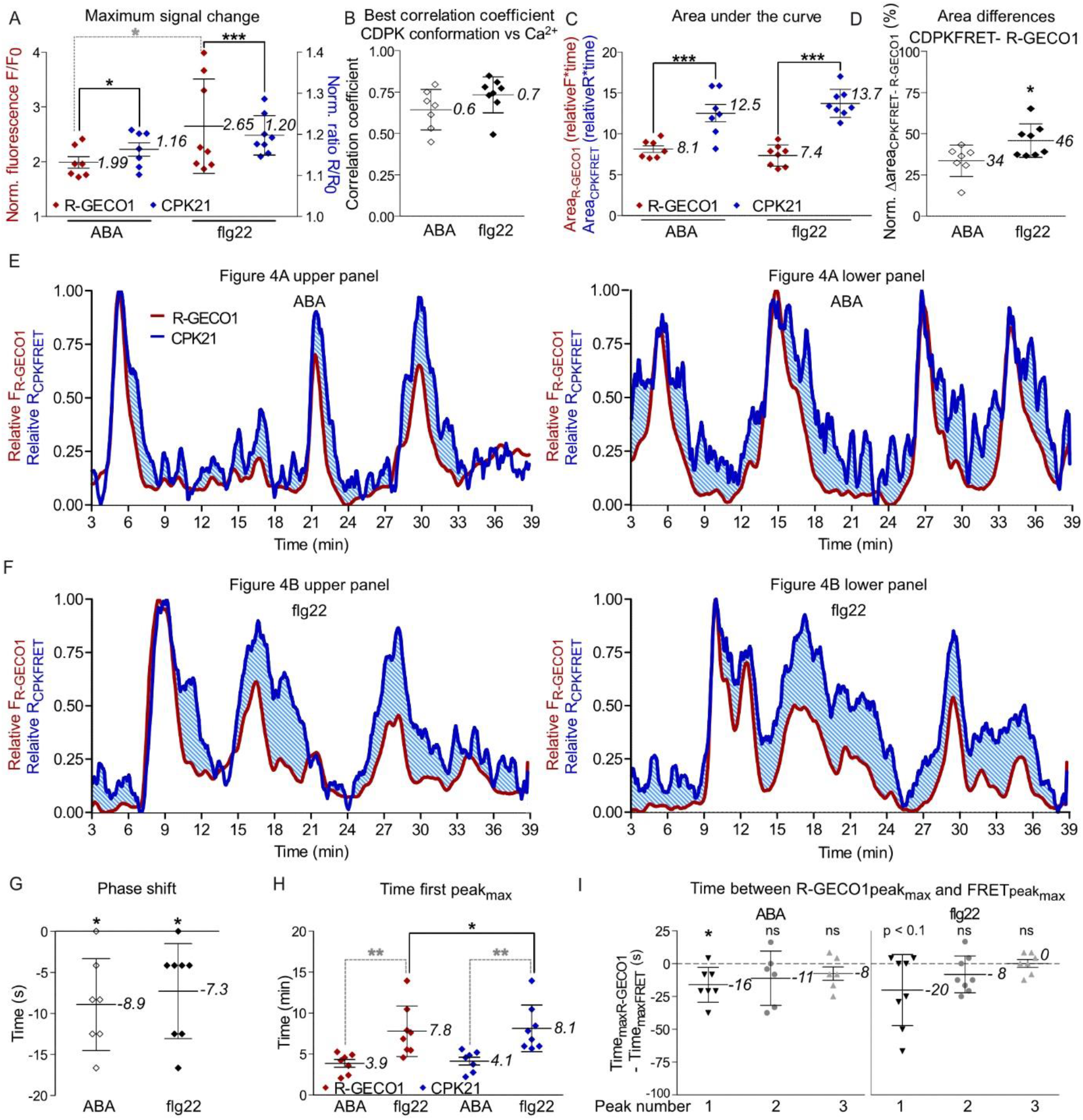
ABA- and flg22-induced CPK21 conformational change signals differ in time, synchronization, strength and shape indicating stimulus-dependent Ca^2+^ signal decoding. A, B, Maximal signal changes (A) and correlation coefficients between R-GECO1 and CPK21-FRET signals (B) after ABA or flg22 application are shown. Adjustment of normalized signal (used in analyses B, E, F) as described in Figure 4. A, B, C, D, G, H, I, Data is represented as the mean ± SD (numbers indicates means), where dots represent the individual measurements (n^ABA^ = 7, n^flg22^ = 8, I, ABA^peak2 and 3^ n= 6, flg22^peak3^ n= 7). Significant differences are indicated by * p ≤ 0.05, ** p ≤ 0.01, *** p ≤ 0.001 and differences with a significant level ≤ 0.1 by ≤ 0.1 or no significant differences (ns). A, Two-way ANOVA and Bonferroni post-test yields for both stimuli significant differences between R-GECO1 and CPK21-FRET, as well as for R-GECO1 between treatments (grey). C-F, Evaluation of decoding by comparative assessment of signal signature (shape of the curves and areas under the curves) from relative signals. For relative signals the signal minimum was set to zero and the maximum to one (E, F). C, Two-way ANOVA and Bonferroni post-test revealed significant differences between R-GECO1 and CPK21-FRET signals. D, By subtracting area^R-GECO1^ from area^CPK21-FRET^ the differences in area are calculated (∆area, light blue shaded area in E, F). ∆ area is normalized to area^CPK21-FRET^. T-test reveals significant differences. Synchronization analysis derived time delays in s (G) and time until first peak maxima in minutes (H). H, Two-way ANOVA and Bonferroni post-test revealed significant differences between treatments (marked in grey), and for flg22 also between R-GECO1 sensor and CPK21-FRET reporter. I, Time differences in s between R-GECO1 and CPK21-FRET ratio-derived first, second and third local peak maxima are shown. G, I, One sided t-test revealed significant differences to zero.

Furthermore, a longer lag time was observed between stimulus application and the appearance of the first Ca^2+^ peak maximum for flg22 (7.8 ± 3.1 min) than for ABA (3.9 ± 1.3 min), and an additional short delay occurred before the first CPK21-FRET recorded peak maximum (Figure 5H). Time differences between R-GECO1 and CDPK-FRET signal phase revealed that cytosolic Ca^2+^ changes precede the CPK21 conformational changes by approximately 8 s for both treatments (Figure 5G). This time delay from Ca^2+^ concentration change to the CPK21 conformational change is most prominent for the first peak, corresponding to an initiation of Ca^2+^ signalling, and decreases during subsequent repetitive Ca^2+^ transients (peak2, peak3) (Figure 5I). Taken together, our data not only identify CPK21 as calcium-sensor kinase that is directly biochemically activated upon ABA- and flg22-treatment in guard cells, but also visualize CPK21-mediated stimulus-specific decoding of distinct Ca^2+^ signatures.

## Discussion

CDPKs are important components of the plant Ca^2+^ regulatory signalling network in response to abiotic and biotic stress and during developmental processes (Boudsocq et al., 2010; Geiger et al., 2010; Geiger et al., 2011; Gutermuth et al., 2013; Matschi et al., 2013; Brandt et al., 2015; Liu et al., 2017; Durian et al., 2020; Fu et al., 2022). One key question is how members of the CDPK gene family are differentially activated and confer distinct signalling function within a single cell. In this study, we developed a FRET-based reporter for the Ca^2+^-induced CDPK conformational change, the first step in CDPK enzyme activation, which precedes ATP-dependent trans-phosphorylation as the second step. Thus, in a biological context CDPK-FRET visualizes the spatial and temporal kinetics of Ca^2+^ decoding as a mean for CDPK activation and function. The CDPK-FRET reporter monitors Ca^2+^-dependent conformational changes of the enzyme within a single molecule (Figure 1). This is in contrast to substrate-based protein kinase FRET reporters, where the substrate protein is sandwiched between a fluorescent protein pair (Brumbaugh et al., 2006; Depry and Zhang, 2011; Zaman et al., 2019; Zhang et al., 2020). CDPKs form a multi-gene family, for which partially redundant *in vivo* functions and overlapping phosphorylations of identical substrate proteins have been reported (Geiger et al., 2010; Geiger et al., 2011; Gutermuth et al., 2013; Brandt et al., 2015; Liu et al., 2017; Gutermuth et al., 2018). Thus, a substrate-based FRET reporter may not yield an isoform-specific resolution for CDPKs *in planta*. By contrast, Ca^2+^-dependent conformational change characterized by K50 and EC50 values are isoform-specific. In addition, regarding the highly conserved modular structure of CDPKs, the here established single molecule CDPK-FRET reporter can be transferred to other CDPK isoforms. Our data identify *in vitro* and *in vivo* for CPK21 a high Ca^2+^-dependency (EC50 = 834 nM) enabling at physiological cytosolic Ca^2+^ levels a significant Ca^2+^-induced conformational change (Figure 3, Figure 4). In comparison, the closest homologue CPK23 follows a different pattern, which includes unique (ABA-dependent) auto-phosphorylation at S362 within its PS domain. The S362D substitution at this site leads from low Ca^2+^-dependency to a constitutive and entirely Ca^2+^-independent activity (Supplemental Figure S3). In *A. thaliana* CPK28, an intrinsic phosphorylation site, S318 within the kinase domain, was shown to influence Ca^2+^-dependency of kinase activity and *in vitro* conformation (Bender et al., 2017; Bredow et al., 2021). These data provide evidence that (auto-) phosphorylation can influence CDPK Ca^2+^-dependency, and both mechanisms may be linked *in planta* to regulate CDPK function in response to biological stimulation.

Stimulus-specific encoding of Ca^2+^ signals and signatures depend on the triggering cue and may be generated at subcellular loci within the cell, determined by the location of stress cue perception and Ca^2+^ influx channels. Thus, appropriate Ca^2+^ decoding is expected within a spatial overlapping region. Interestingly, Ca^2+^ oscillations in the pollen single cell system show a synchronous recording without shifts in phase by both, R-GECO1 and CPK21-FRET. The native *CPK21* gene is not expressed in pollen (Winter et al., 2007) and consistently no biological function for CPK21 in pollen has been reported. Thus, with CPK21 lacking its plasma-membrane binding motifs and present in the cytosol, CPK21-FRET behaves in this CPK21 biologically non-native context as a Ca^2+^ sensor. In guard cells, the unmodified N-terminal domain, important for subcellular localization, guarantees CPK21-specific localization at the plasma membrane (Demir et al., 2013; Simeunovic et al., 2016). In consequence, in *in vivo* context only those fractions of CDPK enzymes are activated in a timely and intracellular spatially distinct manner that can perceive Ca^2+^ changes. CPK21-FRET records ABA- and flg22-induced Ca^2+^ changes in guard cells indicating CPK21 activation, a prerequisite for CPK21 function as Ca^2+^-dependent regulator of anion channel activity. These data not only provide final proof for the so far proposed role of CPK21 as decoder of an ABA-induced Ca^2+^ signature, but also identify CPK21 as candidate for flg22-induced SLAC1 activation. In a biological context, the repetitive transients of Ca^2+^ concentration changes and CPK21 conformational change signal curves differ between both stimuli with a higher signal change dynamic (‘area under the curve’) for flg22 than for ABA (Figure 5C-F). Remarkably, a fairly high synchronization is observed for the Ca^2+^ increase phase, corresponding to rapid Ca^2+^-induced CDPK activation for both stimuli. However, in particular in response to flg22, enzyme reset in conformation and inactivation is delayed compared to the fast decrease for Ca^2+^. Thus, in native guard cells CPK21-FRET reports Ca^2+^ decoding, with the Ca^2+^ signal preceding the CPK21 conformational change during the entire time period of recording. In contrast, in the CPK21 non-native pollen system, the change of Ca^2+^ concentration and CPK21-conformational change are highly synchronous and CDPK-FRET monitors the oscillatory Ca^2+^ pattern similar to the Ca^2+^ sensor R-GECO1.

We interpret the stimulus-specific delay in CDPK-FRET reset upon Ca^2+^ decrease in a functional biological context as a mean to assess decoding. We have demonstrated that post-translational modifications such as (auto-) phosphorylation affects CDPK conformation and activity (Figure 2, Supplemental Figure S3). Also, CDPK may change its assembly within other protein complexes or membrane sub-domains, possibly leading to differences in shape and synchronisation between Ca^2+^ signature and CPK21 decoding. For example, a stress-dependent delocalisation of CPK21 in plasma membrane nanodomains coinciding with CPK21-SLAH3 interaction has been reported (Demir et al., 2013). Furthermore, the differences in response times and synchronisation between cytosolic Ca^2+^ (as reported by R-GECO1) and conformational signals at the plasma membrane (as detected by CDPK FRET) may reflect the spatial separation and kinetics of extracellular Ca^2+^ influx and subsequent sequestration into Ca^2+^ internal stores.

Interestingly, a time delay for the maximal signal change in repetitive Ca^2+^ transients between R-GECO1 and CPK21-FRET signals is most prominent for the first transient (Figure 5I) and diminishes during subsequent phases, indicating that ongoing Ca^2+^ decoding may rely on an in part pre-activated Ca^2+^ decoding machinery.

Our work presents CDPK-FRET as a novel approach to visualize real-time Ca^2+^ signalling and decoding in plant cells. With the increase in number of characterized ion channel types and non-canonical membrane-permeable proteins of overlapping function in Ca^2+^ influx, a shift in the mechanistic understanding of response specificity on the level of Ca^2+^ decoding is in order. Based on isoform-characteristic Ca^2+^-induced conformational change mirroring biochemical activation, CDPK-FRET links in spatial-temporal resolution the decoding of Ca^2+^ signatures to uncover yet unknown function of CDPKs in signalling pathways of interest.

## Methods

### Mutagenesis and cloning of CPK21 and CPK23 enzyme variants

Expression vector *pGEX-6P1* (GE Healthcare) –based recombinant synthesis of *CPK21* and *CPK23* constructs carrying a C-terminal polyhistidine-tag and N-terminal GST-tag has been described previously (Geiger et al., 2010). *CPK21* and *CPK23* in *pGEX-6P1* were used as a template for PCR-based site-directed mutagenesis with primers for the variants CPK21I373S, CPK21I373D, CPK23S362I, CPK23S362D (for primer sequence see Supplemental Table S1) (Weiner et al., 1994). *CPK23CLD21-S362I* chimeric construct was generated by replacing a part of CPK23 pseudosubstrate segment (from aa 353), the CLD23 and a part of *pGEX-6P1* vector backbone in *pGEX-6P1-CPK23* with the homolog sequence from *pGEX-6P1-CPK21* via HindIII and PstI. To create the *pGEX-6P1-CPK23CLD21* construct amino acid substitution I362S was introduced by site-specific mutagenesis primers CPK21I373S-F and CPK21I373S-R with *CPK23CLD21-S362I* in *pGEX-6P1* as PCR template. The *pXCS-CPK23*-HA-StrepII *in vivo* expression plasmid used in this study has been described (Geiger et al., 2010). Mutation of D193 in CPK23 and D204 in CPK21 lead to kinase deficient variants. To create CPK23*D193A* mutagenesis primers CPK23D193A-F and CPK23D193A-R introduced the amino acid substitution D193A.

### Generation of CDPK FRET sensors

*CPK21* and *CPK23* variable and inactive kinase domain coding sequences were re-amplified from plasmids *pXCS-CPK21D204A*-HA-StrepII (Geiger et al., 2011) and *pXCS-CPK23D193A*-HA-StrepII using the forward primer CPK21-VK XbaI EcoRI-F or CPK23-VK XbaI EcoRI-F and the reverse primer CPK21/23-VK XbaI-R. Fragments were transferred via the N- and C-terminal introduced XbaI site into pUC-F3-II (Waadt et al., 2014) resulting in *pUC-F3-II-CPK21*-VK (variable and kinase domain) or *pUC-F3-II*-*CPK23*-VK. *PUC-F3-II* contains the FRET donor (mTurquoise, mT) (Goedhart et al., 2010) and the FRET acceptor (Venus circularly permutated at amino acid 173, cpV173) (Nagai et al., 2004). If not stated otherwise, with CPKk for active kinase, kinase-deficient variants were used for all CDPK-FRET fusion proteins. *CPK21* pseudosubstrate segment and CLD were isolated by PCR, using for linker variant F1 the primers CPK21-PSCLD ApaI-F and CPK21-PSCLD SmaI-R, for linker variant F2 CPK21-PSCLD SpeI-F and CPK21-PSCLD KpnI-R, or for linker variant F3 CPK21-PSCLD BamHI-F and CPK21-PSCLD SalI-R and inserted between ApaI/SmaI (F1), SpeI/KpnI (F2) and BamHI/SalI (F3) in *pUC-F3-II-CPK21*-VK resulting into *pUC-F3-II-CPK21*-FRET (F1-F3), respectively. *CPK21* and *CPK23* coding sequence covering the pseudosubstrate segment and CLD domain until EF-hand 4 (CPK21 aa 522, CPK23 aa 511) was amplified with primers introducing ApaI (CPK21-PSCLD ApaI-F and CPK23-PSCLD ApaI-F) and SmaI (CPK21-PSCLD-EF4 SmaI-R or and CPK23-PSCLD-EF4 SmaI-R) restriction sites. The fragments were inserted between ApaI/SmaI in *pUC-F3-II-CPK21*-VK to yield *pUC-F3-II-CPK21*-FRET (F4), or in *pUC-F3-II-CPK23*-VK to yield *pUC-F3-II-CPK23*-FRET. CPK21 aa 523 and CPK23 aa 512 was identical to the first aa of SmaI restriction site thus in *pUC-F3-II-CPK21*-FRET (F4), or in *pUC-F3-II-CPK23*-FRET CDPK aa sequence until aa 523 (CPK21) or 512 (CPK23) is present. *Escherichia coli* expression vectors were obtained by sub-cloning *CPK21*-FRET (F1-F4) and *CPK23*-FRET via EcoRI/SacI into pET-30a (+) (Novagen). The resulting *pET-30a-CPK21*-FRET (F1-F4) and *pET-30a-CPK23*-FRET constructs carry a N-terminal polyhistidine-tag derived from *pET-30a* vector backbone and a C-terminal StrepII-tag derived from *pUC-F3-II-CPK21-*FRET (F1-F4) or *pUC-F3-II-CPK23*-FRET constructs. For cloning of *CPK21*- and *CPK23*-FRET variants, mutation-containing *CDPK* sequences or the *CPK23CLD2*1 chimeric sequence were exchanged via restriction sites from *pGEX-6P-1-CDPK* constructs into *pET30a-CDPK*-FRET (F4). For *in vivo* transient expression *CPK21*-FRET and *CPK23*-FRET were sub-cloned via EcoRI/Ecl136II and inserted between EcoRI/SfoI into *pXCS-HA-StrepII* (Witte et al., 2004) yielding *pXCS*-*CPK21*- or *pXCS*-*CPK23*-FRET. The *p35S* of *pXCS-*CDPK-FRET was substituted with the *ubiquitin4-2* promoter from parsley from *V69-pUbi:Cas9-MCS-U6* (Kirchner et al., 2017) via AscI/XhoI yielding *pXC-Ubi-CPK21*-FRET or *pXC*-*Ubi-CPK23*-FRET. For cytosolic localization *pXC-Ubi-CDPK*-FRET construct was used as a template for PCR-based site-directed mutagenesis introducing G2A-C3V mutation. To increase the brightness of the FRET donor mT was substituted by eCFP (cyan fluorescent protein) without altering the kinetics of conformational change (Supplemental figure S1E). The primers CFP NdeI-F and CFP ApaI-R were used to amplify *CFP* (Heim and Griesbeck, 2004) with attached NdeI/ApaI sites and *CFP* was cloned into *pXC-Ubi-CPK21G2A-C3V*-FRET and *pXC-Ubi-CPK23G2A-C3V*-FRET or for expression in *E. coli* in *pET-30a-CPK21*-FRET. In all *in vivo* measurements CDPK-FRET fusion constructs with eCFP were used.

For estradiol inducible expression *CPK21*-FRET was cloned in *pER10* this estradiol inducible system for use in transgenic plants has been described elsewhere (Sudarshana et al., 2006). *CPK21*-FRET coding sequences were re-amplified from plasmids *pET-30a-CPK21*-FRET (with eCFP) using the forward primer CPK21 XhoI-F and the reverse primer StrepII SpeI-R and cloned into XhoI SpeI linearized *pER10* yielding *pER10-CPK21*-FRET.

### Generation of transgenic Plants

Transgenic *A. thaliana* line was generated via transforming R-GECO1 (Waadt et al., 2017) plants with p*ER10-CPK21-*FRET by floral dip method. One week old kanamycin resistance seedlings were then transferred to 1 mL distilled water containing 10 µM 17-β-estradiol and 0.05 % DMSO and incubated for 48 hours and screened for fluorescence using a fluorescence stereo zoom microscope (Zeiss Axio Zoom.V16, Zeiss). All selected CPK21-FRET expressing lines showed a patchy expression pattern or unevenly distributed fluorescence as reported previously in estradiol inducible systems (Zuo et al., 2000; Schlücking et al., 2013).

### Expression in *Escherichia coli* and protein purification

CDPK constructs were expressed as recombinant double-tagged fusion proteins in *E. coli*. For *in vitro* kinase assays, expression vector *pGEX-6P-1* was used and proteins were purified using the N-terminal His-Tag and a C-terminal GST-Tag. For *in vitro* FRET measurements expression vector *pET30a*-*CDPK*-FRET was used and proteins were purified using the N-terminal His-Tag and a C-terminal StrepII-Tag.

To synthesize and purify proteins for kinase assays the *pGEX-6P-1*-*CDPK* expression vectors were introduced in *E. coli* BL21 (DE3) (Stratagene). Bacteria were grown at 37 °C in LB medium containing 100 µg/ml ampicillin and protein expression was induced at an OD_600_ of 0.4 - 0.6 with 0.3 mM isopropylthiol-β-galactoside (IPTG) at 28 °C for 4h. Cells were lysed in 4 ml histidine-lysis buffer (50 mM HEPES-KOH pH 7.4, 300 mM NaCl, 0.2 % (v/v) Triton X-100, 1 mM DTT, 10 µl protease inhibitor cocktail for histidine tagged proteins (Sigma) / 0.2 g weight of *E. coli* cells and 30 mM imidazole) using 1*°*mg/ml lysozyme and sonification. After centrifugation supernatant was rotated with 300 – 600 µl Ni sepharose 6 fast flow (GE Healthcare) at 4*°*C for 1 h. Sample/Ni sepharose mix was loaded on empty columns and washed 1 × 10 ml histidine-washing buffer (50 mM HEPES-KOH pH 7.4, 300 mM NaCl) with 30 mM imidazole and 1 × 10 ml histidine-washing buffer with 40 mM imidazole. Proteins were eluted 3 x in 500 µl histidine-elution buffer (50 mM HEPES-KOH pH 7.4, 300 mM NaCl, 500 mM imidazole). Eluate was incubated at 4 °C for 1 h with Glutathione sepharose. Eluate/Glutathione sepharose mix was loaded on columns washed 3 × 3 ml GST-wash buffer (50 mM Tris-HCl pH 8.0, 250 mM NaCl, 1 mM DTT) and eluted 3 x with 300 µl GST-elution buffer (100 mM Tris-HCl pH 8.4 and 20 mM glutathione). Proteins were dialyzed using micro dialysis capsule QuixSep (Roth) and dialysis membrane with 6000 – 8000 Da cut off (Roth). Dialysis-buffer was composed of 30 mM MOPS pH 7.4 and 150 mM KCl.

To synthesize and purify proteins for FRET measurements *pET30a*-*CDPK*-FRET expression vectors were transformed into *E. coli* BL21 (DE3) pLySs strain (Stratagene). Bacteria were grown at 37 °C in TB medium containing 50 µg/ml kanamycin and 34 µg/ml chloramphenicol and protein expression was induced at an OD_600_ of 0.4 - 0.6 with 0.4 mM IPTG at 22 °C for 4h. Cell lyses and purification of histidine tagged proteins as descripted for GST-CDPK-His fusion proteins (see above). His-eluate was incubated at 4°C for 45 min with Strep-tactin macroprep (IBA). StrepII-tagged recombinant proteins were purified as described by Schmidt and Skerra (Schmidt and Skerra, 2007) with the modification that EDTA was omitted from elution and wash buffer. Proteins were dialyzed using micro dialysis capsule QuixSep (Roth) and dialysis membrane with 6000 – 8000 Da cut off (Roth). Dialysis buffer was composed 30 mM Tris-HCl pH 7.4, 150 mM NaCl and 10 mM MgCl_2_. 10 % SDS–PAGE and Coomassie staining confirmed protein purity of *E. coli* expressed proteins. For *in vitro* analyses, protein concentrations were quantified based on the method of Bradford (Protein assay, Bio-Rad).

### Protein sequence comparison

The analysis of the pseudosubstrate segment of the entire *A. thaliana* Col-0 CDPK gene family was conducted on basis of protein sequences from uniprot (Consortium, 2021) with the program Web Logo (Schneider and Stephens, 1990; Crooks et al., 2004).

### Preparation of calcium and magnesium buffers

For CDPK protein kinase assays calculated reciprocal dilutions of zero-Ca^2+^-buffer (10 mM EGTA 150 mM KCl, 30 mM MOPS pH 7.4) with high-Ca^2+^-buffer (10 mM CaCl_2_, 10 mM EGTA 150 mM KCl, 30 mM MOPS pH 7.4) were mixed. For the analysis of CDPK-FRET conformational changes high-Ca^2+^-buffer (20 mM CaCl_2_; 20 mM EGTA, 150 mM NaCl, 10 mM MgCl_2_, 30 mM Tris-HCl pH 7.4) and zero-Ca^2+^-buffer (20 mM EGTA, 150 mM NaCl, 10 mM MgCl_2_, 30 mM Tris-HCl pH 7.4) were mixed accordingly, preceding a 1:1 dilution with CPKs. Correspondingly, buffer solutions for CDPK-FRET analysis in a Mg^2+^ concentration gradient were prepared by a mixture of high-Mg^2+^- buffer (120 mM MgCl_2_, 20 mM EDTA, 150 mM NaCl, 30 mM Tris-HCl pH 7.4) and zero-Mg^2+^-buffer (20 mM EDTA, 150 mM NaCl,, 30 mM Tris-HCl pH 7.4), followed by a 1:1 dilution with FRET protein in Mg^2+^-dialysis buffer (30 mM Tris-HCl pH 7.4, 150 mM NaCl) before data acquisition. The indicated free Ca^2+^- or Mg^2+^-concentrations were calculated with the WEBMXC extended website http://tinyurl.com/y48t33xq based on (Patton et al., 2004).

### *In vitro* kinase assays

*In vitro* kinase activity with recombinant purified proteins were conducted as described using a 20 aa peptide (41-RGPNRGKQRPFRGFSRQVSL-60; JPT Peptide Technologies) derived from the CPK21 and CPK23 *in vivo* phosphorylation substrate protein SLAC1 (slow anion channel-associated 1). For kinase reaction (30 µl) the enzyme (∼90 nM) was incubated in 25 mM MOPS pH 7.4, 125 mM KCl, 10 mM MgCl_2_, 10 µM ATP, 3 μCi [γ-^32^P]-ATP, 10 µM SLAC1 peptide, 6.67 mM EGTA and different concentration of CaCl_2_ (2 - 4 technical replicates per Ca^2+^-concentration) for 20 min at 22°C. The reaction was stopped by adding 3 µl 10 % phosphoric acid. Phosphorylation of the SLAC1 peptide was assessed after binding of phosphor-peptides to P81 filter paper and scintillation counting as described (Franz et al., 2011). Ca^2+^-dependent kinase activity is analysed by a four-parameter logistic equation and indicated as percentage of maximal activity. For analysis of autophosphorylation activities of protein kinases, the reaction is the same as above with the modifications that SLAC1 substrate peptide was omitted from kinase reaction and ∼255 nM enzyme was used. The reaction was stopped by adding 5 × SDS-PAGE loading buffer and boiling for 5 min and samples were separated by 10 % SDS-PAGE. Phosphorylation was determined by autoradiography and phospho-imaging (Typhoon FLA 9500, GE Healthcare).

### *In vitro* analyses of CDPK-FRET

CDPK-FRET protein (∼415 nM) in dialysis buffer diluted 1:1 with Ca^2+^-buffers of defined concentrations was evaluated using the TECAN Infinite M200 PRO plate reader (TECAN). Excitation at 435 nm (bandwidth 5 nm) and emission within the range of 470 - 600 nm was monitored in 2 nm steps with 10 flashes of 20 µs and 400 Hz. cpVenus173/mTurquoise FRET ratios were calculated based on maximal values from emission bands of mTurquoise (470 – 490 nm) and cpVenus173 (518 – 538 nm). FRET ratios are plotted against increasing Ca^2+^-concentrations using a four-parameter logistic equation. The best fit-value obtained for bottom FRET ratio is used to calculate ∆FRET as the percentage change of emission ratios. ∆FRET is defined as:

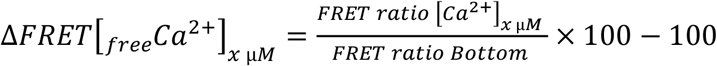

The percentage change of emission ratio ∆FRET is fitted by a four-parameter logistic equation.

The pH-dependency of a CDPK-FRET sensor was assessed with CPK21-FRET in dialysis buffers (30 mM Tris-HCl, 150 mM NaCl and 10 mM MgCl_2_) of different pH values (pH 5.0, 6.9 and 8.0). Dialysed proteins were diluted 1:1 with either high-Ca^2+^- buffer or zero-Ca^2+^-buffer (see above) within 30 mM Tris-HCl adjusted accordingly to a pH range between 5.0 - 8.4.

### Protein expression in protoplast and purification for MS measurement

Preparation and transfection of *Arabidopsis* leaf mesophyll protoplasts from *cpk23* (SALK_007958) (Ma and Wu, 2007) for transient expression of CPK23-HA-StrepII and kinase inactive CPK23D193A-HA-StrepII was conducted as described (Wu et al., 2009). After incubation for 14°h half of the protoplast sample were treated with 30 µM final ABA for 10 min at RT. Cells were collected by centrifugation (twice 10,000 x g, 2 s) and frozen in liquid nitrogen. Protoplasts were resuspended into 600 µl of extraction buffer (100 mM Tris-HCl pH 8.0, 100 mM NaCl, 5 mM EDTA, 5 mM EGTA, 20 mM DTT, 10 mM NaF, 10 mM NaVO_4_, 10 mM β-glycerole-phosphate, 0.5 mM AEBSF, 2 µg/ml aprotinin, 2 µg/ml leupeptin, 100 μg/ml avidin, 0.2 % NP-40, 1 × phosphatase inhibitor mixture (Merck) and 1 × protease inhibitor mixture (Merck)) and centrifuged at 20,000 x g for 10 min at 4 °C. Supernatant was incubated with 24 µl Strep-Tactin MacroPrep (IBA) beads for 45 min at 4 °C. After centrifugation (500 x g, 1 min) beads were solved in 100 µl 6 M urea, 2 M thiourea, pH 8.0 and incubated for 10 min at 4 °C. After centrifugation (500 x g, 1 min) the protein containing supernatant was transferred to a new tube and reduction of disulfide bonds, alkylation of cysteines and tryptic digestion was conducted as described (Dubiella et al., 2013). Peptide containing reactions were vacuum-dried at 30 °C and stored at – 20 °C.

### Targeted analysis phosphorylation by directed MS

Samples were subsequently desalted through C18 tips. Digested protein mixtures were spiked with 500 fmol of 13C_6_-R/K mass-labelled standard peptide before mass spectrometry analysis. Tryptic peptide mixtures including the stable-isotope labelled standard peptides were analysed on a nano-HPLC (Easy nLC, Thermo Scientific) coupled to an Orbitrap mass spectrometer (LTQ-Orbitrap, Thermo Scientific) as mass analyser. Peptides were eluted from a 75 µm analytical column (Easy Columns, Thermo Scientific) on a linear gradient running from 10 % to 30 % acetonitrile in 120 min and were ionized by electrospray. The target peptide V(pS)AVSLSEEEIK (m/z of doubly-charged ion for phosphopeptide 685.8262; non-phosphopeptide 645.8430) was analysed in its phosphorylated and non-phosphorylated state using the stable-isotope labelled synthetic standard peptide as an internal reference and for normalisation between samples. Standards carried a ^13^C_6_-labeled amino acid (arginine or lysine) at their C-terminal ends. Information-dependent acquisition of fragmentation spectra for multiple-charged peptides was used with preferred precursor selection of the target peptides through implementation of an inclusion lists (Schmidt et al., 2011). Full scans were obtained at a resolution of FWHM (full width at half maximum) of 60000, CID fragment spectra were acquired in the LTQ. Additional fragmentation though multistage activation (Schroeder et al., 2004) was used if peptides displayed a loss of phosphoric acid (neutral loss, 98 Da) upon MS/MS fragmentation. Protein identification and intensity quantitation was performed as described (Menz et al., 2016). To allow robust identification and quantitation of the internal standard peptide, multiplicity was set to 2 and Lys6 and Arg6 were selected as stable isotope labels and in general, data analysis was focused on the target peptide sequences only.

### Quantification of target peptide abundance changes

For quantitative analysis, the ion intensities of ^13^C_6_-labeled standard peptides were used for normalisation between samples and replicates. Normalised ion intensities of phosphorylated and non-phosphorylated target peptides were averaged between replicates of the same treatments.

### Transient pollen transformation

*Nicotiana tabacum* (cultivars Petit Havana SR1) plants were grown on soil with a day/night regime of 10 h/14 h, and a temperature of 22 to 24/20 to 22°C provided by a 30 klx white light (SON-T Agro 400W; Philips). Pollen of tobacco lines expressing the R-GECO1 Ca^2+^ sensor (Zhao et al., 2011) as a stable transgene under the control of a pollen specific promoter (*pLeLAT52::R-GECO1* line) was used from frozen stocks to perform transient transformation using a homemade particle bombardment device recently described in detail (Gutermuth et al., 2013). Biolistic transformation was performed with *pXC-Ubi-CPK21-G2A-C3V-CFP* and *pXC-Ubi-CPK23-G2A-C3V-CFP* on agar plates containing pollen tube growth medium (1 mM MES-Tris pH 5.8, 0.2 mM CaCl_2_, 9.6 mM HCl, and 1.6 mM H_3_BO_3_). Osmolality of pollen media was adjusted to 400 mosmol kg^-1^ (Vapor Pressure Osmometer 5520) with D(+)-sucrose. Tip-focused dynamic Ca^2+^ oscillation patterns were provoked by using a medium containing 10 mM Cl^-^.

### Live-cell fluorescence imaging in pollen tubes

The setup for wide-field live-cell imaging and the appropriate software to control sample acquisition has been described in detail (Gutermuth et al., 2013). Images were recorded with a time interval of 5 s. For simultaneous CFP/YFP/RFP-imaging a triple-band dichroic mirror (Chroma # 69008; ET - ECFP/EYFP/mCherry) was used to reflect excitation light on the samples. Excitation of CFP and R-GECO1 was performed with a VisiChrome High-Speed Polychromator System (Visitron Systems) at 420 nm and 550 nm, respectively. Optical filters (Chroma Technology Corporation) for CFP (ET 470/24 nm), YFP (ET 535/30 nm) and R-GECO1 (624/40) were used for fluorescence detection with a back-illuminated 512 × 512 pixel Evolve EMCCD camera (Photometrics). A high-speed 6-position filter wheel (Ludl Electronic Products Ltd.) ensured the quasi simultaneous imaging of all three channels with a lag-time of ∼0.1 sec. For image processing the following steps were conducted for R-GECO1 (R-GECO1_excitation_/R-GECO1_emission_), FRET (CFP_excitation_/YFP_emission_) and CFP (CFP_excitation_/CFP_emission_) channels using Fiji (National Institute of Health) (Schindelin et al., 2012) background subtraction (same value for FRET and CFP channel), gaussian blur, 32-bit conversion, kymographs were generated, and threshold adjusted. A self-made script for the Octave 4.0.3 free software http://www.gnu.org/software/octave/) was used to quantify fluorescence intensities of each channel at ∼5 - 15 µm behind the tip of the growing pollen tubes over time as described in (Gutermuth et al., 2018). FRET-analysis was performed by dividing the FRET signal by the CFP signal. Synchronization analyses were performed with R v.4.1 (R Core Team, 2022) as described in live-cell fluorescence imaging in guard cell.

### Live-cell fluorescence imaging in guard cell

For epidermal peal sample preparation leaf material from 2 – 3 week old plants grown on jiffy-7 soil (Jiffy Products) under short-day conditions (10 h day light, 20 – 22°C; 60 % RH) were used. Epidermal peel sample preparation in 2 well chambered coverslips (IBIDI) has been described in detail (Eichstädt et al., 2021). Fresh prepared epidermis single-layer was directly immersed with 1 ml plant buffer (10 mM MES-Tris pH 6.15, 5 mM KCl, 50 μM CaCl_2_, 20 µM 17-β-estradiol and 0.05 % DMSO) and the samples were incubated for recovery overnight. The samples were incubated under light at 20–22°C for at least 2 h before imaging. Confocal imaging was performed in bottom imaging mode on a Zeiss LSM 880 system (Zeiss) with a 40 × water immersion objective (LD C-APOCHROME, 40 x/1.1 Korr UV-VIS M27; Zeiss). 16-bit images were acquired every 4.16 s with a frame size of 512 × 512 pixels and a pinhole of 599 µm. Fluorescence proteins were excited with 458 (eCFP_excitation_/eCFP _emission_ and eCFP_excitation_/YFP_emission_), 514 (YFP_excitation_/YFP_emission_) and 561 (R-GECO1_excitation_/R-GECO1_emission_) nm and an emission-range between 465 and 505 nm for eCFP, 525 and 560 nm for YFP or 580 and 611 nm for R-GECO1 was used for detection. Guard cells showing a constant resting Ca^2+^ level were selected to analyze Ca^2+^ increase upon stimulus application. The nature and function of spontaneous Ca^2+^ transients occurring in some guard cells were discussed elsewhere (Allen et al., 1999; Klüsener et al., 2002; Hubbard et al., 2012). For 100 nM flg22 and 20 µM ABA treatments, 50-fold concentrations were prepared in water (flg22) or 10 % ethanol (ABA) and added in a 1:50 volume ratio. Treatment with ethanol (0.2 % final) as solvent control did not induce measurable signal increases of R-GECO1 or CPK21-FRET (Supplemental Figure S8). For image processing the following steps were conducted for R-GECO1 (R-GECO1_excitation_/R-GECO1_emission_), FRET (CFP_excitation_/YFP_emission_) and CFP (CFP_excitation_/CFP_emission_) channels using Fiji: gaussian blur filter set to 1, 32-bit conversion, threshold adjustment (stack histogram) and selection of one guard cell as region of interest (ROI). The mean gray values of these ROIs were used for further calculations. The FRET ratio was calculated via dividing FRET (CFP_excitation_/YFP_emission_) mean gray values over CFP (CFP_excitation_/CFP_emission_) mean gray values. The resulting FRET emission ratio and the R-GECO1 signal was normalized to the mean of the 10 frames before treatment (R/R_0_) or (F/F_0_). For graphic representation intensity-over-time plots data were plotted and smoothed (averaging 15 values on each side using a second order polynomial) with GraphPad Prism 5 software. Before synchronisation analysis of stomata measurements, the normalized FRET emission ratio and normalized R-GECO1 fluorescence were adjusted to signal changes derived from technical artefacts (like stomata movement) by calculating a trend line. All following steps were performed in R v. 4.1 with R studio (R Core Team, 2022; Team, 2022) using the R packages *scales* (H. Wickham et al., 2022a), *readxl* (H. Wickham et al., 2022b), *writexl* (Ooms, 2021) *stats* and *forecast* (R. J Hyndman and Khandakar, 2008; Hyndman R et al., 2022). For calculating this trend line, a linear regression model was applied with time as predictor variable and signal (normalized FRET emission ratio and normalized R-GECO1 fluorescence) as response variable assuming that the technical artefacts can be represented by a linear relationship. The values of the linear regression model were used to revise the data by dividing the normalized signals by the corresponding data of the linear regression. Synchronization analysis for Ca^2+^ and FRET conformational change were used to quantify their phase relationship by a cross correlation analysis with an R-script called phase_analysis which was described previously (Li et al., 2021). To analyse the signal change dynamics (area under the curve) the following steps were conducted in R v. 4.1: the revised data were smooth by a moving average (8 data points; centred) and rescaled between 0 (minimum of the signals) and 1 (maximum of the signals) for the time points after treatment application. The area under curve data was calculated using GraphPad Prism 5.

### Statistics

Analyses of kinase activities and conformational changes were performed by a four-parameter logistic equation with a global model with shared hillslope for all data sets of the same enzyme using GraphPad Prism 5. For comparison of enzyme variants data from different measurements were combined in one figure. Statistical analyses of variance (t-Test, ANOVA) were performed using GraphPad Prism 5.

## Supporting information

Suppemental figure S1-S9, Supplemental table S1

## Accession numbers

CPK21 (AT4G04720) and CPK23 (AT4G04740)

## Supplemental data

Supplemental Figure S1: Design and characterization of CDPK-FRET reporter

Supplemental Figure S2: Kinase activity measurements CPK21 conformational change sensor

Supplemental Figure S3: Characterization of CPK23 auto-phosphorylation

Supplemental Figure S4: Tip-focused intracellular Ca^2+^ gradients and CPK21 conformational change upon transient expression of CPK21G2A-C3V FRET fusion protein in tobacco pollen tubes.

Supplemental Figure S5: CPK23 is unable to decode the tip-focused intracellular Ca^2+^ gradient and Ca^2+^-oscillations in growing tobacco pollen tubes.

Supplemental Figure S6: ABA dependent Ca^2+^ concentration transients in guard cells are decoded by CPK21.

Supplemental Figure S7: CPK21-FRET decodes flg22-induced cytosolic Ca^2+^- concentration transients in guard cells.

Supplemental Figure S8: Flg22 but not the solvent control EtOH induce cytosolic Ca^2+^ concentration transients and CPK21 conformational changes.

Supplemental Figure S9 Synchronization analysis between CPK21-FRET and R-GECO1 in pollen tube tip.

Supplemental Movie S1 Tip-focused intracellular Ca^2+^ gradient and Ca^2+^ oscillations in a growing tobacco pollen expressing CPK21G2A-C3V-FRET.

Supplemental Movie S2 Tip-focused intracellular Ca^2+^ gradient and Ca^2+^ oscillations in a growing tobacco pollen expressing CPK23G2A-C3V-FRET

Supplemental Table S1: Sequences of oligonucleotide primers

## Acknowledgments

We thank Tim-Martin Ehnert, and Sylvia Krüger for technical assistance and Klara Altintoprak for contributing to the mutagenesis and cloning of CPK21 and CPK23 enzyme variants, and Simon Gilroy, Helle Ulrich and Roman Lassig for critical comments on the manuscript. We would like to acknowledge the assistance of the FU Berlin Core Facility BioSupraMol supported by the Deutsche Forschungsgemeinschaft (DFG). This work was funded within DFG grant (Ko3657/2-3) to K.R.K. and DFG research unit FOR964 and collaborative research centre CRC973 to T.R..

## Contributions

A.L. conceived the study and established conformational change analysis using CDPK-FRET, performed all *in vitro* kinase assays and conducted *in vivo* imaging. B.E. performed the cloning of FRET-constructs and all *in vitro* conformational change analyses. S. L. wrote R-scripts and performed analyses; P.S. conducted protein expression and preparation for MS analysis. J. O. conducted the expression analyses of pEstradiol: CPK21-FRET line. W.X.S. performed MS analysis and evaluation. K.R.K. designed and supervised *in vivo* imaging experiments in pollen tubes and generated the tobacco line used. J. A. F. participate in strategic discussions and contribute to experimental design. T.R. conceived the study and contributed to the experimental design. A.L. and T.R. wrote the manuscript. All authors discussed the results and commented on the manuscript.

## Data availability

All data generated in this study are included within the main text and Supplemental information.

## References

Allen, G.J., Kwak, J.M., Chu, S.P., Llopis, J., Tsien, R.Y., Harper, J.F., and Schroeder, J.I. (1999). Cameleon calcium indicator reports cytoplasmic calcium dynamics in Arabidopsis guard cells. The Plant journal : for cell and molecular biology 19, 735–747.

Bender, K.W., Zielinski, R.E., and Huber, S.C. (2018). Revisiting paradigms of Ca2+ signaling protein kinase regulation in plants. Biochem. J. 475, 207–223.

Bender, K.W., Blackburn, R.K., Monaghan, J., Derbyshire, P., Menke, F.L., Zipfel, C., Goshe, M.B., Zielinski, R.E., and Huber, S.C. (2017). Autophosphorylation-based Calcium (Ca(2+)) Sensitivity Priming and Ca(2+)/Calmodulin Inhibition of Arabidopsis thaliana Ca(2+)-dependent Protein Kinase 28 (CPK28). The Journal of biological chemistry 292, 3988–4002.

Bi, G., Su, M., Li, N., Liang, Y., Dang, S., Xu, J., Hu, M., Wang, J., Zou, M., Deng, Y., Li, Q., Huang, S., Li, J., Chai, J., He, K., Chen, Y.H., and Zhou, J.M. (2021). The ZAR1 resistosome is a calcium-permeable channel triggering plant immune signaling. Cell 184, 3528–3541.e3512.

Bjornson, M., Pimprikar, P., Nürnberger, T., and Zipfel, C. (2021). The transcriptional landscape of Arabidopsis thaliana pattern-triggered immunity. Nature plants 7, 579–586.

Boudsocq, M., Droillard, M.-J., Regad, L., and Laurière, C. (2012). Characterization of Arabidopsis calcium-dependent protein kinases: activated or not by calcium? The Biochemical journal 447, 291–299.

Boudsocq, M., Willmann, M.R., McCormack, M., Lee, H., Shan, L., He, P., Bush, J., Cheng, S.-H., and Sheen, J. (2010). Differential innate immune signalling via Ca2+ sensor protein kinases. Nature 464, 418–422.

Brandt, B., Munemasa, S., Wang, C., Nguyen, D., Yong, T., Yang, P.G., Poretsky, E., Belknap, T.F., Waadt, R., Alemán, F., and Schroeder, J.I. (2015). Calcium specificity signaling mechanisms in abscisic acid signal transduction in Arabidopsis guard cells. eLife 4.

Bredow, M., Bender, K.W., Johnson Dingee, A., Holmes, D.R., Thomson, A., Ciren, D., Tanney, C.A.S., Dunning, K.E., Trujillo, M., Huber, S.C., and Monaghan, J. (2021). Phosphorylation-dependent subfunctionalization of the calcium-dependent protein kinase CPK28. Proceedings of the National Academy of Sciences of the United States of America 118.

Brumbaugh, J., Schleifenbaum, A., Gasch, A., Sattler, M., and Schultz, C. (2006). A dual parameter FRET probe for measuring PKC and PKA activity in living cells. Journal of the American Chemical Society 128, 24–25.

Consortium, U. (2021). UniProt: the universal protein knowledgebase in 2021. Nucleic acids research 49, D480–d489.

Crooks, G.E., Hon, G., Chandonia, J.-M., and Brenner, S.E. (2004). WebLogo: A sequence logo generator. Genome Res 14, 1188–1190.

Damineli, D.S.C., Portes, M.T., and Feijó, J.A. (2017). Oscillatory signatures underlie growth regimes in Arabidopsis pollen tubes: computational methods to estimate tip location, periodicity, and synchronization in growing cells. Journal of experimental botany 68, 3267–3281.

Demir, F., Horntrich, C., Blachutzik, J.O., Scherzer, S., Reinders, Y., Kierszniowska, S., Schulze, W.X., Harms, G.S., Hedrich, R., Geiger, D., and Kreuzer, I. (2013). Arabidopsis nanodomain-delimited ABA signaling pathway regulates the anion channel SLAH3. Proceedings of the National Academy of Sciences of the United States of America 110, 8296–8301.

Depry, C., and Zhang, J. (2011). Using FRET-based reporters to visualize subcellular dynamics of protein kinase A activity. Methods in molecular biology 756, 285–294.

Dubiella, U., Seybold, H., Durian, G., Komander, E., Lassig, R., Witte, C.-P., Schulze, W.X., and Romeis, T. (2013). Calcium-dependent protein kinase/NADPH oxidase activation circuit is required for rapid defense signal propagation. Proceedings of the National Academy of Sciences 110, 8744–8749.

Durian, G., Sedaghatmehr, M., Matallana-Ramirez, L.P., Schilling, S.M., Schaepe, S., Guerra, T., Herde, M., Witte, C.P., Mueller-Roeber, B., Schulze, W.X., Balazadeh, S., and Romeis, T. (2020). Calcium-dependent protein kinase CPK1 controls cell death by in vivo phosphorylation of senescence master regulator ORE1. The Plant cell 32, 1610–1625.

Eichstädt, B., Lederer, S., Trempel, F., Jiang, X., Guerra, T., Waadt, R., Lee, J., Liese, A., and Romeis, T. (2021). Plant Immune Memory in Systemic Tissue Does Not Involve Changes in Rapid Calcium Signaling. Frontiers in plant science 12, 798230.

Felle, H.H. (2001). pH: Signal and Messenger in Plant Cells 3, 577–591.

Franz, S., Ehlert, B., Liese, A., Kurth, J., Cazalé, A.-C., and Romeis, T. (2011). Calcium-dependent protein kinase CPK21 functions in abiotic stress response in Arabidopsis thaliana. Molecular plant 4, 83–96.

Fu, D., Zhang, Z., Wallrad, L., Wang, Z., Höller, S., Ju, C., Schmitz-Thom, I., Huang, P., Wang, L., Peiter, E., Kudla, J., and Wang, C. (2022). Ca(2+)-dependent phosphorylation of NRAMP1 by CPK21 and CPK23 facilitates manganese uptake and homeostasis in Arabidopsis. Proceedings of the National Academy of Sciences of the United States of America 119, e2204574119.

Geiger, D., Scherzer, S., Mumm, P., Marten, I., Ache, P., Matschi, S., Liese, A., Wellmann, C., Al-Rasheid, K.A.S., Grill, E., Romeis, T., and Hedrich, R. (2010). Guard cell anion channel SLAC1 is regulated by CDPK protein kinases with distinct Ca2+ affinities. Proceedings of the National Academy of Sciences of the United States of America 107, 8023–8028.

Geiger, D., Maierhofer, T., AL-Rasheid, K.A.S., Scherzer, S., Mumm, P., Liese, A., Ache, P., Wellmann, C., Marten, I., Grill, E., Romeis, T., and Hedrich, R. (2011). Stomatal closure by fast abscisic acid signaling is mediated by the guard cell anion channel SLAH3 and the receptor RCAR1. Sci Signal 4, ra32..

Gifford, J.L., Walsh, M.P., and Vogel, H.J. (2007). Structures and metal-ion-binding properties of the Ca2+-binding helix–loop–helix EF-hand motifs. The Biochemical journal 405, 199–221.

Goedhart, J., van Weeren, L., Hink, M.A., Vischer, N.O., Jalink, K., and Gadella, T.W., Jr. (2010). Bright cyan fluorescent protein variants identified by fluorescence lifetime screening. Nature methods 7, 137–139.

Guo, J., He, J., Dehesh, K., Cui, X., and Yang, Z. (2022). CamelliA-based simultaneous imaging of Ca2+ dynamics in subcellular compartments. Plant physiology 188, 2253–2271.

Gutermuth, T., Herbell, S., Lassig, R., Brosché, M., Romeis, T., Feijó, J.A., Hedrich, R., and Konrad, K.R. (2018). Tip-localized Ca(2+)-permeable channels control pollen tube growth via kinase-dependent R- and S-type anion channel regulation. The New phytologist 218, 1089–1105.

Gutermuth, T., Lassig, R., Portes, M.-T., Maierhofer, T., Romeis, T., Borst, J.-W., Hedrich, R., Feijó, J.A., and Konrad, K.R. (2013). Pollen tube growth regulation by free anions depends on the interaction between the anion channel SLAH3 and calcium-dependent protein kinases CPK2 and CPK20. The Plant cell 25, 4525–4543.

Guzel Deger, A., Scherzer, S., Nuhkat, M., Kedzierska, J., Kollist, H., Brosché, M., Unyayar, S., Boudsocq, M., Hedrich, R., and Roelfsema, M.R. (2015). Guard cell SLAC1-type anion channels mediate flagellin-induced stomatal closure. The New phytologist 208, 162–173.

H. Wickham, D. Seidel, and RStudio. (2022a). Graphical scales map data to aesthetics, and provide methods for automatically determining breaks and labels for axes and legends.

H. Wickham, Bryan, J., and RStudio. (2022b). readxl: Read Excel Files.

Harmon, A.C., Gribskov, M., and Harper, J.F. (2000). CDPKs - a kinase for every Ca2+ signal? Trends Plant Sci 5, 154–159.

Heim, N., and Griesbeck, O. (2004). Genetically encoded indicators of cellular calcium dynamics based on troponin C and green fluorescent protein. The Journal of biological chemistry 279, 14280–14286.

Huang, J.-Z., Hardin, S.C., and Huber, S.C. (2001). Identification of a novel phosphorylation motif for CDPKs: Phosphorylation of synthetic peptides lacking basic residues at P-3/P-4. Arch. Biochem. Biophys. 393, 61–66.

Huang, S., Waadt, R., Nuhkat, M., Kollist, H., Hedrich, R., and Roelfsema, M.R.G. (2019). Calcium signals in guard cells enhance the efficiency by which abscisic acid triggers stomatal closure. New Phytol 224, 177–187.

Hubbard, K.E., Siegel, R.S., Valerio, G., Brandt, B., and Schroeder, J.I. (2012). Abscisic acid and CO2 signalling via calcium sensitivity priming in guard cells, new CDPK mutant phenotypes and a method for improved resolution of stomatal stimulus– response analyses. Ann. of Bot. 109, 5–17.

Hyndman R, Athanasopoulos G, Bergmeir C, Caceres G, Chhay L, O’Hara-Wild M, Petropoulos F, Razbash S, Wang E, and f, Y. (2022). forecast: Forecasting functions for time series and linear models.

Keinath, N.F., Waadt, R., Brugman, R., Schroeder, J.I., Grossmann, G., Schumacher, K., and Krebs, M. (2015). Live Cell Imaging with R-GECO1 Sheds Light on flg22- and Chitin-Induced Transient [Ca(2+)]cyt Patterns in Arabidopsis. Molecular plant 8, 1188–1200.

Kirchner, T.W., Niehaus, M., Debener, T., Schenk, M.K., and Herde, M. (2017). Efficient generation of mutations mediated by CRISPR/Cas9 in the hairy root transformation system of Brassica carinata. PloS one 12, e0185429.

Klüsener, B., Young, J.J., Murata, Y., Allen, G.J., Mori, I.C., Hugouvieux, V., and Schroeder, J.I. (2002). Convergence of calcium signaling pathways of pathogenic elicitors and abscisic acid in Arabidopsis guard cells. Plant Physiol 130, 2152–2163.

Konrad, K.R., Wudick, M.M., and Feijó, J.A. (2011). Calcium regulation of tip growth: new genes for old mechanisms. Current opinion in plant biology 14, 721–730.

Köster, P., DeFalco, T.A., and Zipfel, C. (2022). Ca(2+) signals in plant immunity. The EMBO journal 41, e110741.

Kudla, J., Becker, D., Grill, E., Hedrich, R., Hippler, M., Kummer, U., Parniske, M., Romeis, T., and Schumacher, K. (2018). Advances and current challenges in calcium signaling. New Phytol. 218, 414–431.

Li, K., Prada, J., Damineli, D.S.C., Liese, A., Romeis, T., Dandekar, T., Feijó, J.A., Hedrich, R., and Konrad, K.R. (2021). An optimized genetically encoded dual reporter for simultaneous ratio imaging of Ca(2+) and H(+) reveals new insights into ion signaling in plants. The New phytologist 230, 2292–2310.

Liese, A., and Romeis, T. (2013). Biochemical regulation of in vivo function of plant calcium-dependent protein kinases (CDPK). Biochim Biophys Acta 1833, 1582–1589.

Liu, K.H., Niu, Y., Konishi, M., Wu, Y., Du, H., Sun Chung, H., Li, L., Boudsocq, M., McCormack, M., Maekawa, S., Ishida, T., Zhang, C., Shokat, K., Yanagisawa, S., and Sheen, J. (2017). Discovery of nitrate-CPK-NLP signalling in central nutrient-growth networks. Nature 545, 311–316.

Ma, S.-Y., and Wu, W.-H. (2007). AtCPK23 functions in Arabidopsis responses to drought and salt stresses. Plant Mol Biol 65, 511–518.

Matschi, S., Werner, S., Schulze, W.X., Legen, J., Hilger, H.H., and Romeis, T. (2013). Function of calcium-dependent protein kinase CPK28 of Arabidopsis thaliana in plant stem elongation and vascular development. The Plant journal : for cell and molecular biology 73, 883–896.

Menz, J., Li, Z., Schulze, W.X., and Ludewig, U. (2016). Early nitrogen-deprivation responses in Arabidopsis roots reveal distinct differences on transcriptome and (phospho-) proteome levels between nitrate and ammonium nutrition. The Plant journal : for cell and molecular biology 88, 717–734.

Michard, E., Dias, P., and Feijó, J.A. (2008). Tobacco pollen tubes as cellular models for ion dynamics: improved spatial and temporal resolution of extracellular flux and free cytosolic concentration of calcium and protons using pHluorin and YC3.1 CaMeleon. Sex Plant Reprod 21, 169–181.

Michard, E., Simon, A.A., Tavares, B., Wudick, M.M., and Feijó, J.A. (2017). Signaling with Ions: The Keystone for Apical Cell Growth and Morphogenesis in Pollen Tubes. Plant physiology 173, 91–111.

Mou, W., Kao, Y.T., Michard, E., Simon, A.A., Li, D., Wudick, M.M., Lizzio, M.A., Feijó, J.A., and Chang, C. (2020). Ethylene-independent signaling by the ethylene precursor ACC in Arabidopsis ovular pollen tube attraction. Nature communications 11, 4082.

Nagai, T., Yamada, S., Tominaga, T., Ichikawa, M., and Miyawaki, A. (2004). Expanded dynamic range of fluorescent indicators for Ca(2+) by circularly permuted yellow fluorescent proteins. Proceedings of the National Academy of Sciences of the United States of America 101, 10554–10559.

Ojo, K.K., Larson, E.T., Keyloun, K.R., Castaneda, L.J., DeRocher, A.E., Inampudi, K.K., Kim, J.E., Arakaki, T.L., Murphy, R.C., Zhang, L., Napuli, A.J., Maly, D.J., Verlinde, C.L.M.J., Buckner, F.S., Parsons, M., Hol, W.G.J., Merritt, E.A., and Van Voorhis, W.C. (2010). Toxoplasma gondii calcium-dependent protein kinase 1 is a target for selective kinase inhibitors. Nat Struct Mol Biol 17, 602–607.

Ooms, J. (2021). Zero-dependency data frame to xlsx exporter based on ‘libxlsxwriter’. Fast and no Java or Excel required.

Patton, C., Thompson, S., and Epel, D. (2004). Some precautions in using chelators to buffer metals in biological solutions. Cell calcium 35, 427–431.

R Core Team. (2022). R: A language and environment for statistical computing 1.

R. J Hyndman, and Khandakar, Y. (2008). Automatic Time Series Forecasting: the forecast Package for R. Journal of Statistical Software 27.

Saris, N.E., Mervaala, E., Karppanen, H., Khawaja, J.A., and Lewenstam, A. (2000). Magnesium. An update on physiological, clinical and analytical aspects. Clinica chimica acta; international journal of clinical chemistry 294, 1–26.

Scherzer, S., Maierhofer, T., Al-Rasheid, K.A.S., Geiger, D., and Hedrich, R. (2012). Multiple calcium-dependent kinases modulate ABA-activated guard cell anion channels. Mol Plant. 5, 1409–1412.

Schindelin, J., Arganda-Carreras, I., Frise, E., Kaynig, V., Longair, M., Pietzsch, T., Preibisch, S., Rueden, C., Saalfeld, S., Schmid, B., Tinevez, J.Y., White, D.J., Hartenstein, V., Eliceiri, K., Tomancak, P., and Cardona, A. (2012). Fiji: an open-source platform for biological-image analysis. Nature methods 9, 676–682.

Schlücking, K., Edel, K.H., Köster, P., Drerup, M.M., Eckert, C., Steinhorst, L., Waadt, R., Batistic, O., and Kudla, J. (2013). A new β-estradiol-inducible vector set that facilitates easy construction and efficient expression of transgenes reveals CBL3-dependent cytoplasm to tonoplast translocation of CIPK5. Molecular plant 6, 1814–1829.

Schmidt, A., Beck, M., Malmström, J., Lam, H., Claassen, M., Campbell, D., and Aebersold, R. (2011). Absolute quantification of microbial proteomes at different states by directed mass spectrometry. Molecular systems biology 7, 510.

Schmidt, T.G., and Skerra, A. (2007). The Strep-tag system for one-step purification and high-affinity detection or capturing of proteins. Nature protocols 2, 1528–1535.

Schneider, T.D., and Stephens, R.M. (1990). Sequence logos: a new way to display consensus sequences. Nucleic acids research 18, 6097–6100.

Schroeder, M.J., Shabanowitz, J., Schwartz, J.C., Hunt, D.F., and Coon, J.J. (2004). A neutral loss activation method for improved phosphopeptide sequence analysis by quadrupole ion trap mass spectrometry. Analytical chemistry 76, 3590–3598.

Simeunovic, A., Mair, A., Wurzinger, B., and Teige, M. (2016). Know where your clients are: subcellular localization and targets of calcium-dependent protein kinases. Journal of experimental botany 67, 3855–3872.

Sudarshana, M.R., Plesha, M.A., Uratsu, S.L., Falk, B.W., Dandekar, A.M., Huang, T.K., and McDonald, K.A. (2006). A chemically inducible cucumber mosaic virus amplicon system for expression of heterologous proteins in plant tissues. Plant biotechnology journal 4, 551–559.

Tan, Y.Q., Yang, Y., Shen, X., Zhu, M., Shen, J., Zhang, W., Hu, H., and Wang, Y.F. (2022). Multiple cyclic nucleotide-gated channels function as ABA-activated Ca2+ channels required for ABA-induced stomatal closure in Arabidopsis. The Plant cell.

Team, R. (2022). RStudio: Integrated Development Environment for R. RStudio. PBC, Boston, MA 1.

Thor, K., and Peiter, E. (2014). Cytosolic calcium signals elicited by the pathogen-associated molecular pattern flg22 in stomatal guard cells are of an oscillatory nature. The New phytologist 204, 873–881.

Thor, K., Jiang, S., Michard, E., George, J., Scherzer, S., Huang, S., Dindas, J., Derbyshire, P., Leitão, N., DeFalco, T.A., Köster, P., Hunter, K., Kimura, S., Gronnier, J., Stransfeld, L., Kadota, Y., Bücherl, C.A., Charpentier, M., Wrzaczek, M., MacLean, D., Oldroyd, G.E.D., Menke, F.L.H., Roelfsema, M.R.G., Hedrich, R., Feijó, J., and Zipfel, C. (2020). The calcium-permeable channel OSCA1.3 regulates plant stomatal immunity. Nature 585, 569–573.

Tian, W., Wang, C., Gao, Q., Li, L., and Luan, S. (2020). Calcium spikes, waves and oscillations in plant development and biotic interactions. Nature plants 6, 750–759.

Tian, W., Hou, C., Ren, Z., Wang, C., Zhao, F., Dahlbeck, D., Hu, S., Zhang, L., Niu, Q., Li, L., Staskawicz, B.J., and Luan, S. (2019). A calmodulin-gated calcium channel links pathogen patterns to plant immunity. Nature 572, 131–135.

Toyota, M., Spencer, D., Sawai-Toyota, S., Jiaqi, W., Zhang, T., Koo, A.J., Howe, G.A., and Gilroy, S. (2018). Glutamate triggers long-distance, calcium-based plant defense signaling. Science (New York, N.Y.) 361, 1112–1115.

Waadt, R., Krebs, M., Kudla, J., and Schumacher, K. (2017). Multiparameter imaging of calcium and abscisic acid and high-resolution quantitative calcium measurements using R-GECO1-mTurquoise in Arabidopsis. The New phytologist 216, 303–320.

Waadt, R., Hitomi, K., Nishimura, N., Hitomi, C., Adams, S.R., Getzoff, E.D., and Schroeder, J.I. (2014). FRET-based reporters for the direct visualization of abscisic acid concentration changes and distribution in Arabidopsis. eLife 3, e01739.

Waadt, R., Köster, P., Andrés, Z., Waadt, C., Bradamante, G., Lampou, K., Kudla, J., and Schumacher, K. (2020). Dual-Reporting Transcriptionally Linked Genetically Encoded Fluorescent Indicators Resolve the Spatiotemporal Coordination of Cytosolic Abscisic Acid and Second Messenger Dynamics in Arabidopsis. The Plant cell 32, 2582–2601.

Wang, Z., and Gou, X. (2021). The First Line of Defense: Receptor-like Protein Kinase-Mediated Stomatal Immunity. International journal of molecular sciences 23.

Weiner, M.P., Costa, G.L., Schoettlin, W., Cline, J., Mathur, E., and Bauer, J.C. (1994). Site-directed mutagenesis of double-stranded DNA by the polymerase chain reaction. Gene 151, 119–123.

Wernimont, A.K., Artz, J.D., Finerty, P., Lin, Y.-H., Amani, M., Allali-Hassani, A., Senisterra, G., Vedadi, M., Tempel, W., Mackenzie, F., Chau, I., Lourido, S., Sibley, L.D., and Hui, R. (2010). Structures of apicomplexan calcium-dependent protein kinases reveal mechanism of activation by calcium. Nat Struct Mol Biol 17, 596–601.

Winter, D., Vinegar, B., Nahal, H., Ammar, R., Wilson, G.V., and Provart, N.J. (2007). An “Electronic Fluorescent Pictograph” browser for exploring and analyzing large-scale biological data sets. PloS one 2, e718.

Witte, C.P., Noël, L., Gielbert, J., Parker, J., Romeis, T., and (2004). Rapid one-step protein purification from plant material using the eight-amino acid StrepII epitope. Plant Mol Biol 55, 135–147.

Wu, F.-H., Shen, S.-C., Lee, L.-Y., Lee, S.-H., Chan, M.-T., and Lin, C.-S. (2009). Tape-arabidopsis sandwich - a simpler arabidopsis protoplast isolation method. Plant Methods 5, 16.

Xu, G., Moeder, W., Yoshioka, K., and Shan, L. (2022). A tale of many families: calcium channels in plant immunity. The Plant cell 34, 1551–1567.

Yip Delormel, T., and Boudsocq, M. (2019). Properties and functions of calcium-dependent protein kinases and their relatives in Arabidopsis thaliana. The New phytologist 224, 585–604.

Yuan, F., Yang, H., Xue, Y., Kong, D., Ye, R., Li, C., Zhang, J., Theprungsirikul, L., Shrift, T., Krichilsky, B., Johnson, D.M., Swift, G.B., He, Y., Siedow, J.N., and Pei, Z.M. (2014). OSCA1 mediates osmotic-stress-evoked Ca2+ increases vital for osmosensing in Arabidopsis. Nature 514, 367–371.

Zaman, N., Seitz, K., Kabir, M., George-Schreder, L.S., Shepstone, I., Liu, Y., Zhang, S., and Krysan, P.J. (2019). A Förster resonance energy transfer sensor for live-cell imaging of mitogen-activated protein kinase activity in Arabidopsis. The Plant journal : for cell and molecular biology 97, 970–983.

Zhang, L., Takahashi, Y., Hsu, P.K., Kollist, H., Merilo, E., Krysan, P.J., and Schroeder, J.I. (2020). FRET kinase sensor development reveals SnRK2/OST1 activation by ABA but not by MeJA and high CO(2) during stomatal closure. eLife 9.

Zhao, Y., Araki, S., Wu, J., Teramoto, T., Chang, Y.F., Nakano, M., Abdelfattah, A.S., Fujiwara, M., Ishihara, T., Nagai, T., and Campbell, R.E. (2011). An expanded palette of genetically encoded Ca²+ indicators. Science 333, 1888–1891.

Zhou, J.Y., Hao, D.L., and Yang, G.Z. (2021). Regulation of Cytosolic pH: The Contributions of Plant Plasma Membrane H(+)-ATPases and Multiple Transporters. International journal of molecular sciences 22.

Zuo, J., Niu, Q.W., and Chua, N.H. (2000). Technical advance: An estrogen receptor-based transactivator XVE mediates highly inducible gene expression in transgenic plants. The Plant journal : for cell and molecular biology 24, 265–273.

