## Supplementary material for "Imaging of plant calcium-sensor kinase conformation monitors real time calcium decoding *in planta*": Suppemental figure S1-S9, Supplemental table S1

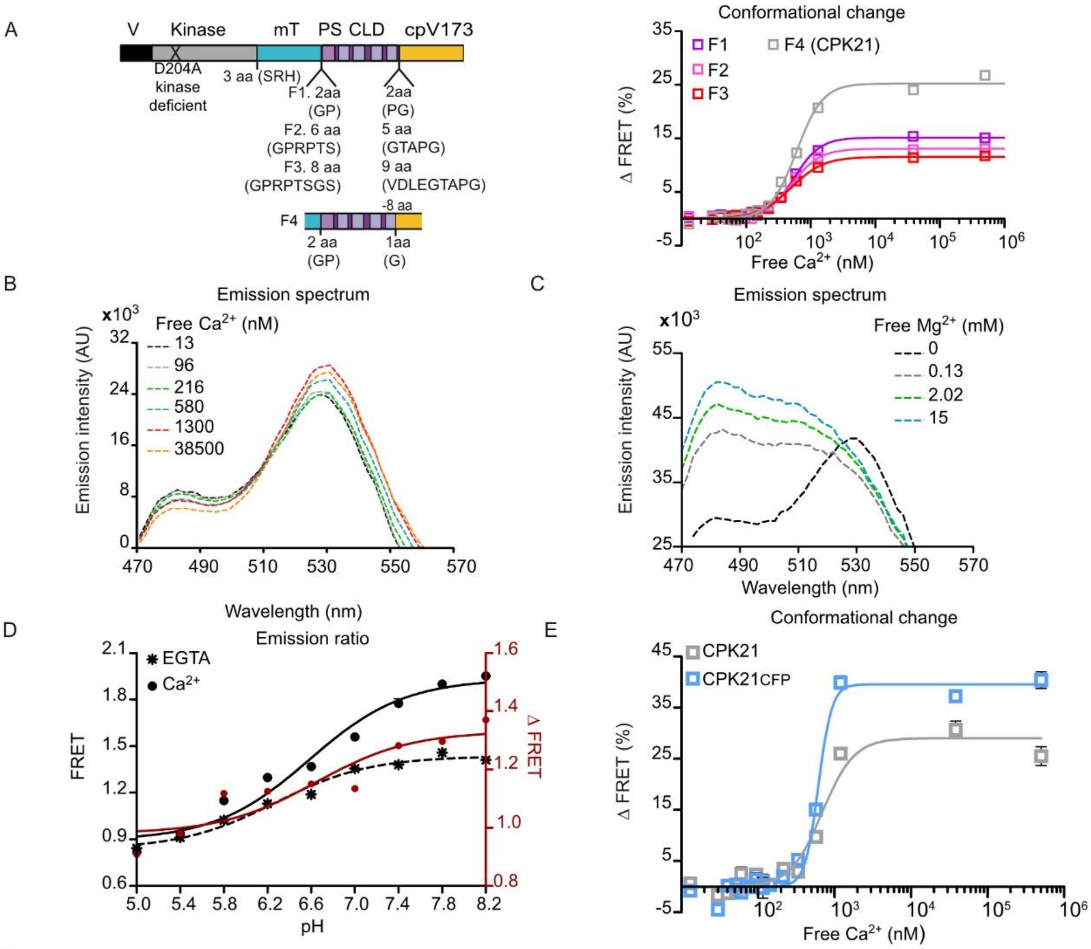


F

| | Conformational change | | | |
|---|---|---|---|---|
| CPK-FRET | EC50 Ca²⁺ (nM) | Δ FRETmax (%) | Hillslope | n |
| CPK21F1 | 571 +/- 9 | 14 +/- 1 | 2.0 | 2 |
| CPK21F2 | 515 +/- 2 | 12 +/- 1 | 2.0 | 2 |
| CPK21F3 | 554 +/- 58 | 10 +/- 1 | 1.7 | 2 |
| CPK21CFP | 708 +/- 89 | 45 +/- 6 | 5.0 | 2 |

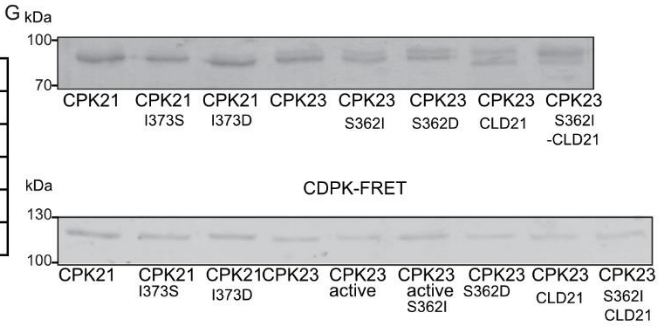


**Supplemental Figure S1**: Design and characterization of CDPK-FRET reporter

A $Ca^{2+}$-dependent conformational change of kinase deficient CPK21 linker variants F1 – F4 (right panel). Numbers indicate the insertion or the deletion of amino acids (aa) with the linker aa sequence shown (left panel). FRET efficiency change is determined as in Fig. 1D ($R^2_{F1}$ = 0.99, $R^2_{F2}$ = 0.98, $R^2_{F3}$ = 0.98, $R^2_{F4}$ = 0.99; mean ± SEM n= 3, 15 $Ca^{2+}$-concentrations). Variant F4 shows the highest change in emission ratio and was used as matrix in all further FRET experiments denominated as CPK21. The CPK21 EC50 value is shown in Fig. 1E. B, C, Emission spectra (470–570 nm with a step size of 1 or 2 nm) of CPK21 at the indicated $Ca^{2+}$ -(B) or $Mg^{2+}$ - (C) concentrations. For $Mg^{2+}$ the experiment was repeated three times with similar results. D, pH-dependency of CPK21 FRET efficiency. FRET-recorded conformational change of CPK21 (left axis) plotted against 9 pH concentrations in the absence of $Ca^{2+}$ (EGTA, mean ± SEM n= 5) and in presence of saturating $Ca^{2+}$-concentration (mean ± SEM n= 5). The ratio FRET ($Ca^{2+}$) / FRET (EGTA) is indicated in red (right axis). The experiment was repeated two times with similar results. E, Comparison of FRET-recorded conformational change of CPK21 with either mTurquoise (CPK21) or CFP (CPK21CFP) as FRET donor ($R^2_{CPK21}$ = 0.94, $R^2_{CPK21CFP}$ = 0.97; mean ± SEM n= 4, 15 $Ca^{2+}$-concentrations). F, Summary of EC50 (conformational change) and maximal FRET efficiency (mean ± SEM) and shared Hillslope in n independent experiments. G, Coomassie stained SDS gel confirms recombinant purified proteins CPK21 and CPK23 variants (top panel; kinase assays) or CPK21-FRET and CPK23-FRET variants (lower panel, conformational change assays).

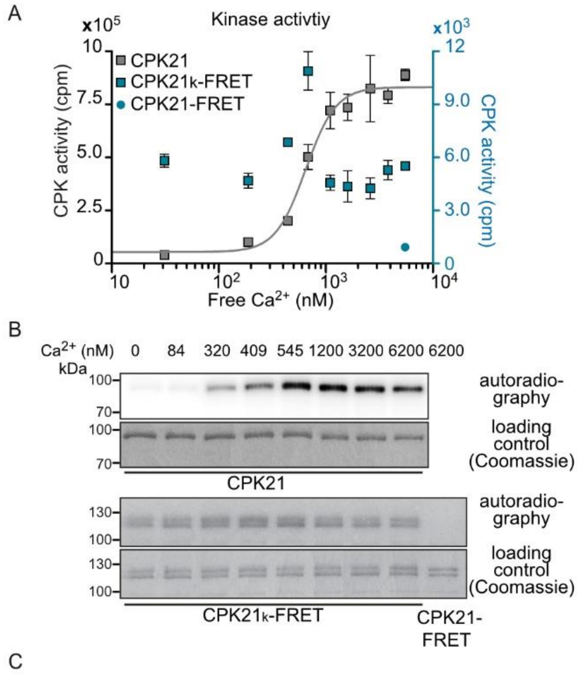


C

| | Kinase activity | | | |
|---|---|---|---|---|
| | $K_{50}$ $Ca^{2+}$ (nM) | core activity (%) | Hillslope | n |
| $CPK21_k$ FRET | not applicable | | | 2 |

**Supplemental Figure S2:** Kinase activity measurements CPK21 conformational change sensor

A, Kinase activity of CPK21 and $CPK21_k$-FRET variant with active kinase domain (plotted against $Ca^{2+}$-concentrations ($R^2_{CPK21}$ = 0.94; mean ± SEM n= 2–3, 9 $Ca^{2+}$-concentrations); for the kinase inactive variant CPK21-FRET (blue dot in Supplemental Figure S2A) only one $Ca^{2+}$-concentration was tested. B, $Ca^{2+}$-dependent auto-phosphorylation of CPK21 and $CPK21_k$-FRET. Kinase reaction mixture was separated by SDS gel electrophoreses followed by autoradiography. The assay was repeated two times with similar results. C, The kinase activity of $CPK21_k$-FRET can`t be analyzed via a Hill equation in n independent experiments.

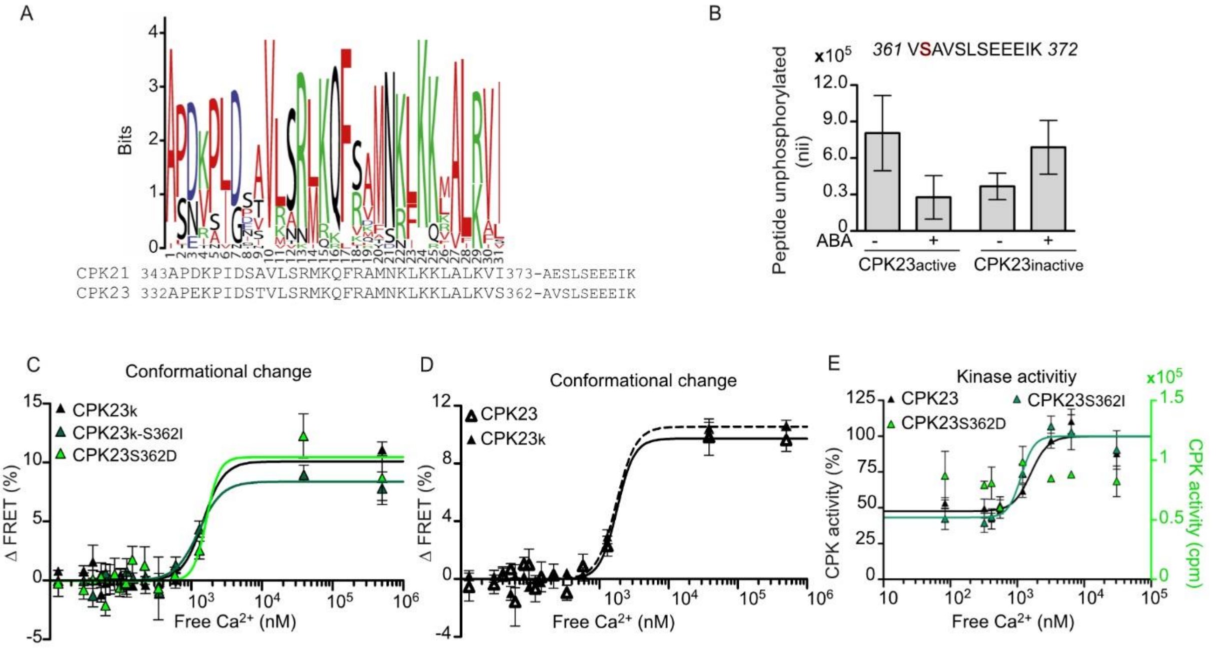


F

| | Conformational change | | | |
|---|---|---|---|---|
| CPK-FRET | EC50 Ca$^{2+}$ (nM) | Δ FRETmax (%) | Hillslope | n |
| CPK23kS362I | 1387 +/- 116 | 9 +/- 1 | 2.6 | 2 |
| CPK23S362D | 1271 +/- 128 | 11 | 4.7 | 4 |
| CPK23k | 1533 +/- 67 | 9 +/- 1 | 3.1 | 5 |

| | Kinase activity | | | |
|---|---|---|---|---|
| CPK | K50 Ca$^{2+}$ (nM) | core activity (%) | Hillslope | n |
| CPK23S362I | 1231 +/- 40 | 37 +/- 6 | 4.9 | 3 |
| CPK23S362D | not applicable | | | 2 |

**Supplemental Figure S3**: Characterization of CPK23 auto-phosphorylation

A, Comparison of PS amino acid sequences of the *A. thaliana* CPK gene family using the program WebLogo with indicated sequences for CPK21 (aa 343 – 373) and CPK23 (aa 332 – 362). B, Protoplasts transiently expressing active CPK23 or kinase inactive CPK23 were treated with ABA, and the amount of unphosphorylated peptide containing S362 (red in sequence above) was quantified via SRM mass spectrometry. Mean ± SEM combining three experiments. C, FRET-recorded conformational change of CPK23k-FRETvariant with active kinase domain, CPK23k-S362I and CPK23S362D plotted against Ca$^{2+}$-concentrations ($R^2_{CPK23k}$ = 0.84, $R^2_{CPK23kS362I}$ = 0.84, $R^2_{CPK23S362D}$ = 0.56). FRET efficiency change is determined as in Fig. 1D (mean ± SEM n= 3-6, 15 Ca$^{2+}$-concentration). D, FRET-recorded conformational change of active CPK23k compared to kinase-inactive CPK23-FRET variant ($R^2_{CPK23}$ = 0.71, $R^2_{CPK23k}$ = 0.91; mean ± SEM n= 5 – 6, 15 Ca$^{2+}$-concentrations). E, Kinase activity of CPK23 PS variants carrying amino acid substitution CPK23S362I and CPK23S362D plotted against Ca$^{2+}$ concentrations ($R^2_{CPK23}$ = 0.84, $R^2_{CPK23S362I}$ = 0.8). Activity is expressed as percentage of full Ca$^{2+}$-saturation (mean ± SEM n= 2-3, 7-8 Ca$^{2+}$-concentration). CPK23S362D activity indicated in counts per minute (cpm) F, Summary of EC50 (conformational change) and K50 (kinase activity) (mean ± SEM) and shared Hillslope in n independent experiments. The kinase activity of CPK23S362D can`t be analyzed via a Hill equation.

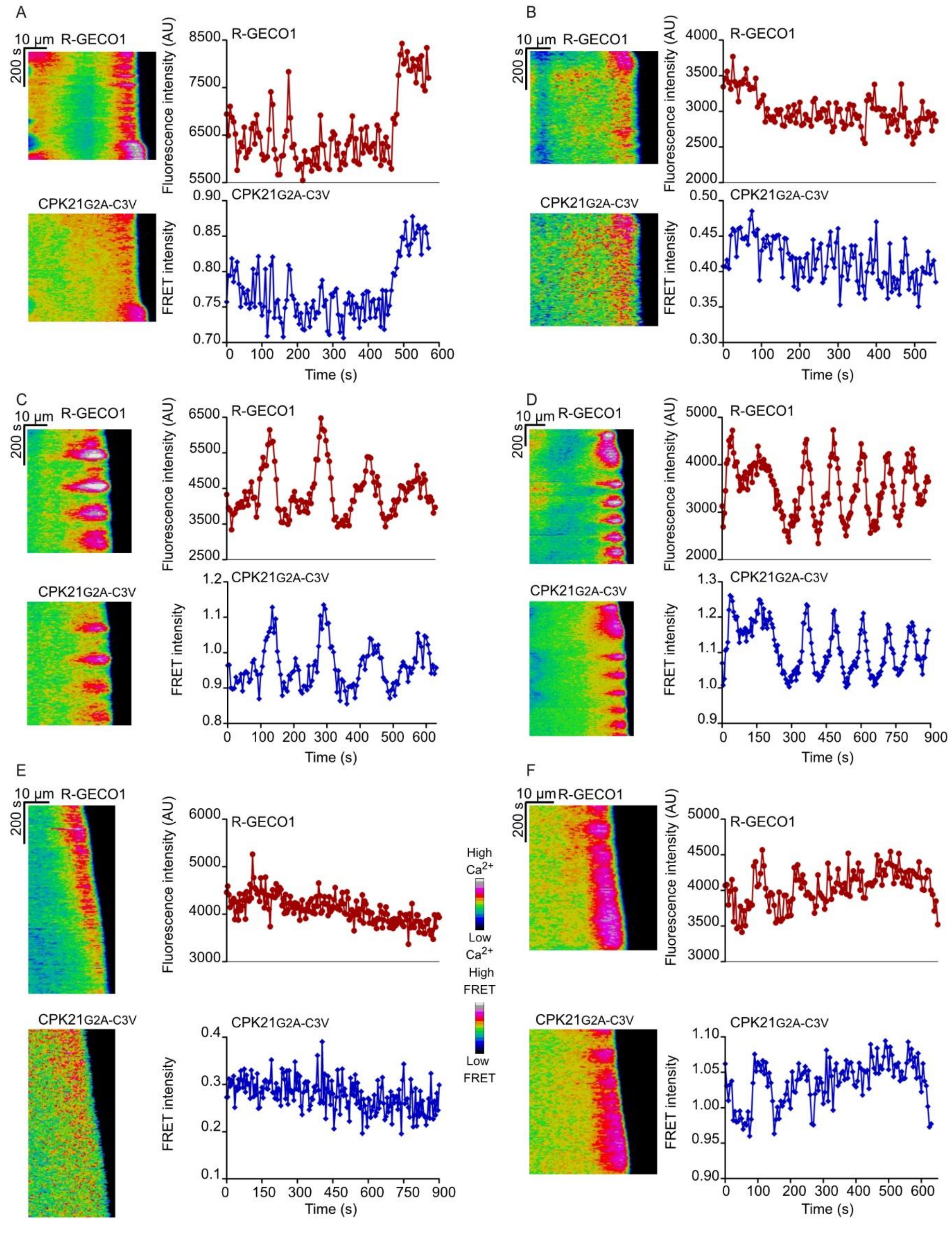


**Supplemental Figure S4**: Tip-focused intracellular $Ca^{2+}$ gradients and CPK21 conformational change upon transient expression of CPK21G2A-C3V FRET fusion protein in tobacco pollen tubes.

Kymographs are deduced from time-lapse imaging of fluorescence intensities by R-GECO1 $Ca^{2+}$ and CPK21-FRET in the tip of pollen tubes. Images were taken every 5 s for 570 s (A), 555 s (B), 630 s (C), 890 s (D), 905 s (E), and 630 s (F).

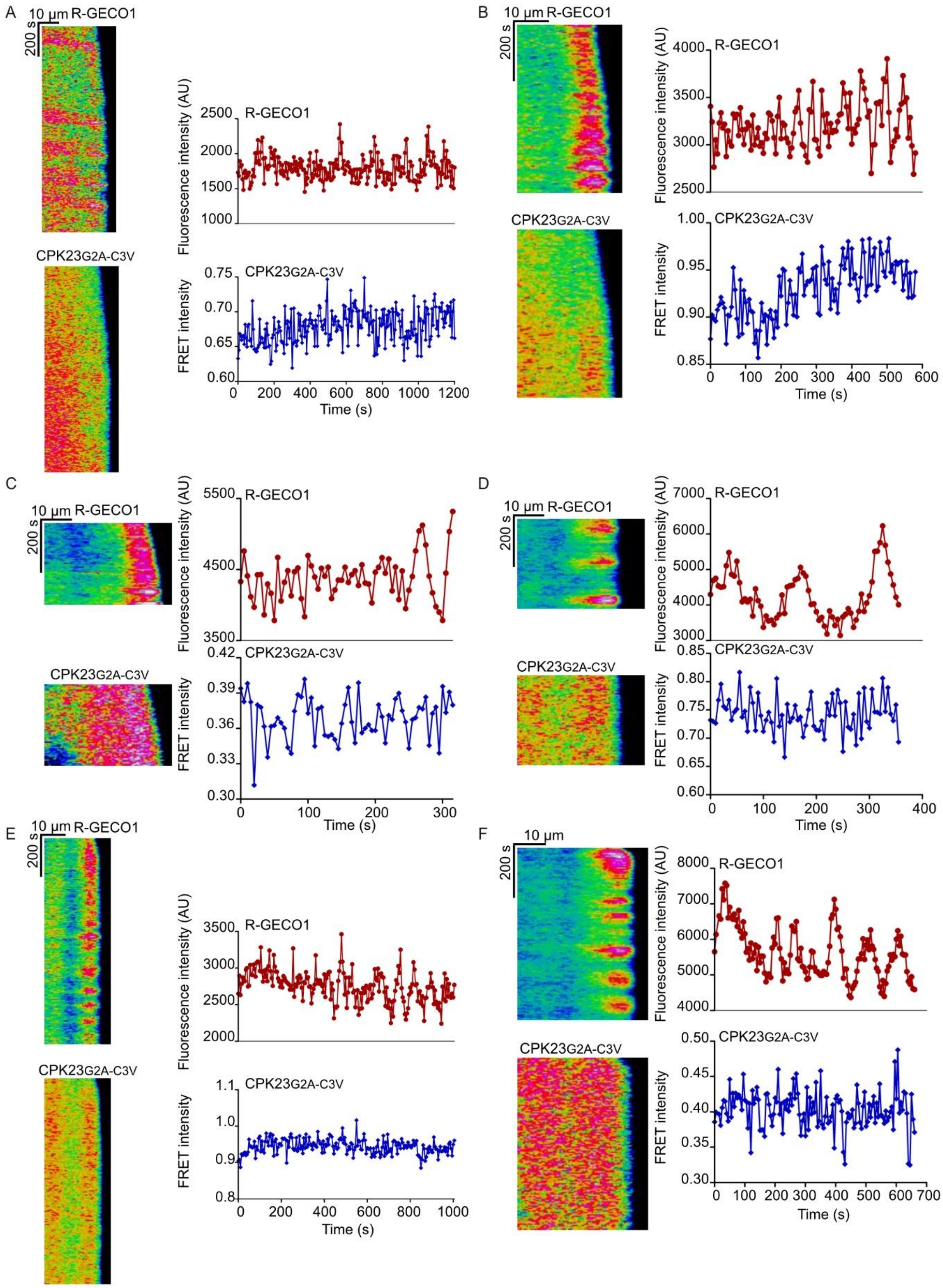


**Supplemental Figure S5**: CPK23 is unable to decode the tip-focused intracellular $Ca^{2+}$ gradient and $Ca^{2+}$-oscillations in growing tobacco pollen tubes.

Time-dependent fluorescence intensity changes were monitored at the tip of the growing pollen tube after transient expression of CPK23 conformation sensor. Images were taken every 5 s for 1205 s (A), 580 s (B), 325 s (C), 355 s (D), 1005 s (E) and 660 s (F).

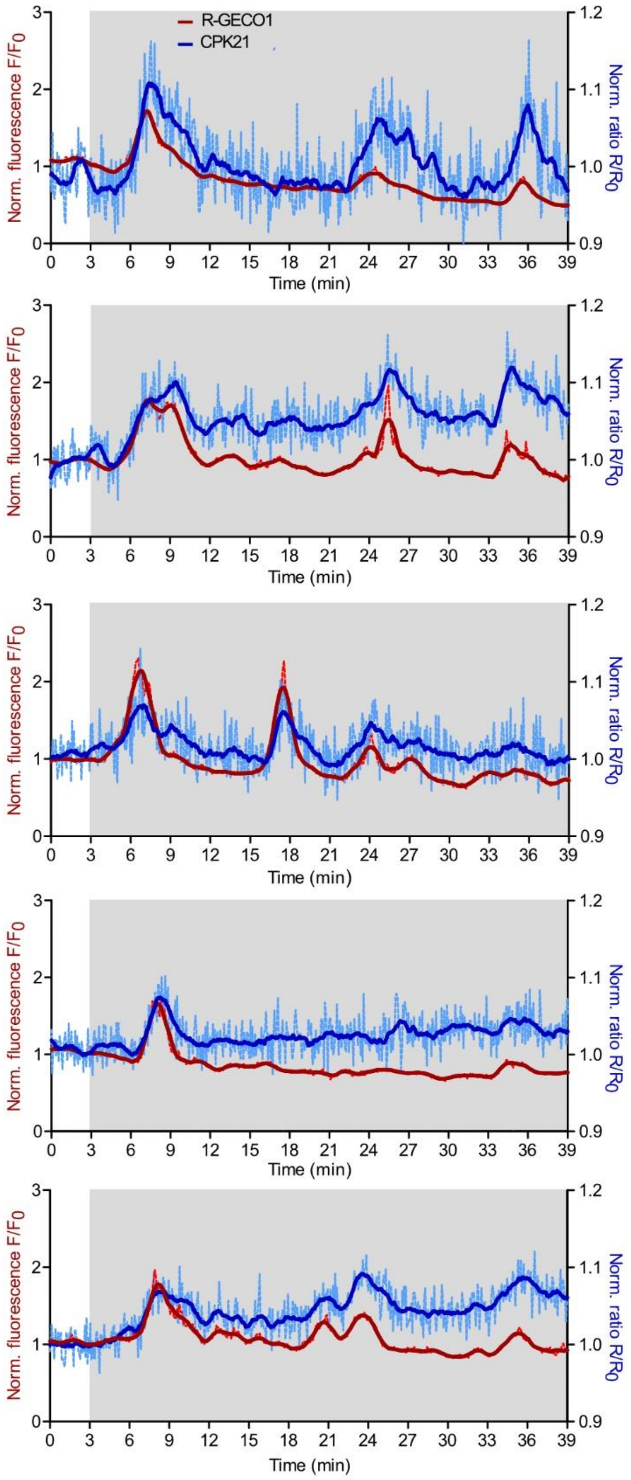


**Supplemental Figure S6**: ABA dependent $Ca^{2+}$ concentration transients in guard cells are decoded by CPK21.

Changes in the cytosolic $Ca^{2+}$ concentration (red curve) and FRET ratio changes (blue curve) were visualized in response to 20 µM ABA. Intensity-over-time plots are shown. Normalized data shown as dotted lighter coloured lines and smooth data (averaging 15 values on each side using a second order polynomial) shown as continuous lines. The ratio (CPK21-FRET) or the fluorescence (R-GECO1) were normalized to the mean value over 10 cycles (around 40 s) before treatment. The time interval of recording after ABA treatment is indicated by the grey area.

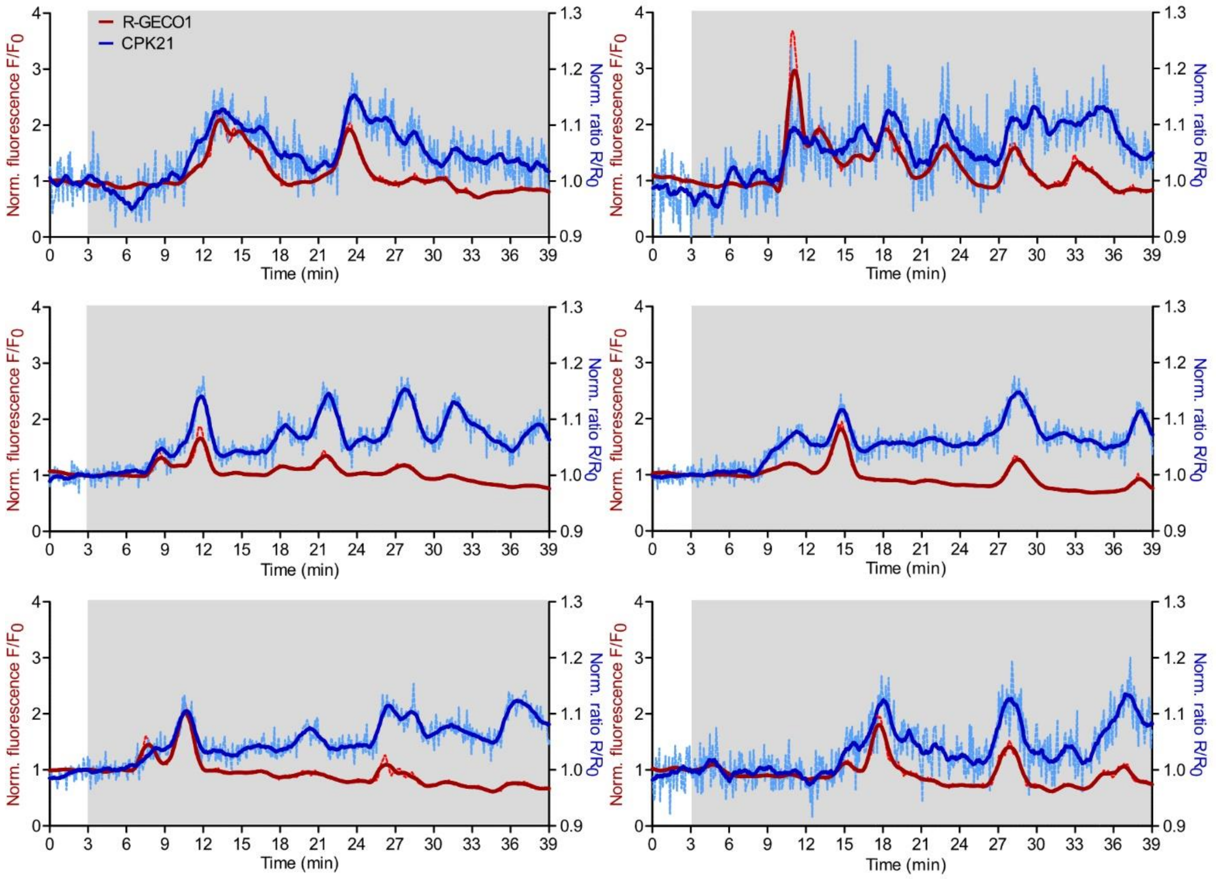


**Supplemental Figure S7**: CPK21-FRET decodes flg22-induced cytosolic $Ca^{2+}$-concentration transients in guard cells.

Time-dependent R-GECO1 fluorescence intensity changes and CPK21-FRET signal changes induced by 100 nM flg22 were monitored in time-lapse *in vivo* imaging. Normalized data shown as dotted lighter coloured lines and smooth data (averaging 15 values on each side using a second order polynomial) shown as continuous lines. flg22 was added at 3 min, the time interval of recording after flg22 treatment is indicated by the grey area. The ratio (CPK21-FRET) or the fluorescence (R-GECO1) were normalized to the mean value over 10 cycles (around 40 s) before treatment.

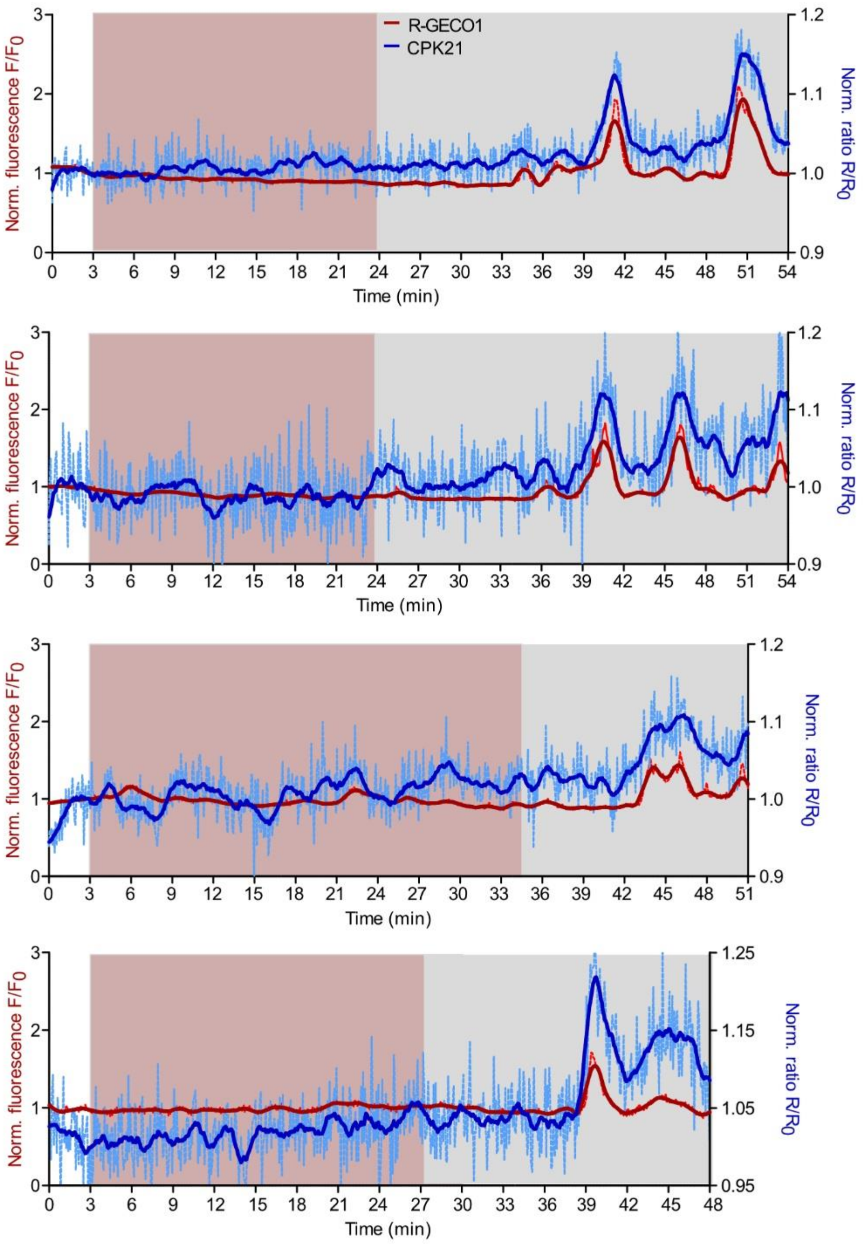


**Supplemental Figure S8**: Flg22 but not the solvent control EtOH induce cytosolic $Ca^{2+}$ concentration transients and CPK21 conformational changes.

In guard cells the cytosolic $Ca^{2+}$ concentration (red curve) and the CPK21-FRET ratio (blue curve) were imaged upon EtOH (0.2 %, solvent control for ABA; magenta area), followed by 100 nM flg22 as second treatment (grey area). Continuous lines in intensity-over-time plots represents smooth data (averaging 15 values on each side using a second order polynomial) and dotted lighter coloured lines shows unsmoothed data. The ratio (CPK21-FRET) or the fluorescence (R-GECO1) was normalized to the mean value over 10 cycles (around 40 s) before treatment.

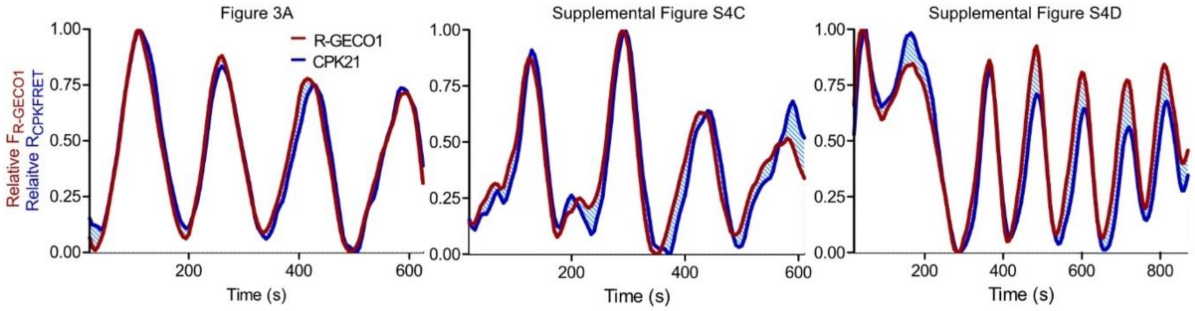


**Supplemental Figure S9** Synchronization analysis between CPK21-FRET and R-GECO1 in pollen tube tip.

To analyse and compare the shape of the $Ca^{2+}$ - and CDPK conformational change signals the minimum was set to zero and the maximum to one (relative signals). Light blue shaded area illustrates the area difference between the $Ca^{2+}$ signal change and CPK21 conformational changes. Numbers above panels indicate corresponding figures containing the original data.

**Supplemental Movie S1** Tip-focused intracellular $Ca^{2+}$ gradient and $Ca^{2+}$ oscillations in a growing tobacco pollen expressing CPK21G2A-C3V-FRET. Tobacco pollen carrying the R-GECO1 $Ca^{2+}$-sensor as a stable transgene were transiently transformed with CDPK-FRET conformational change sensors containing CPK21G2A-C3V-FRET. Time-dependent fluorescence intensity changes induced by a medium containing 10 mM $Cl^{-}$ were recorded as time-laps video. The time-laps video represents the same pollen tube as shown in Figure 3A. The images were taken every 5 s for 650 s, assembled as a stack and converted into a video (10 frames/s, 1 s in the video equals to 50 s in the recording).

**Supplemental Movie S2** Tip-focused intracellular $Ca^{2+}$ gradient and $Ca^{2+}$ oscillations in a growing tobacco pollen expressing CPK23G2A-C3V- FRET. Tobacco pollen carrying the R-GECO1 $Ca^{2+}$-sensor as a stable transgene was transiently transformed with CPK23G2A-C3V-FRET. Tip-focused dynamic $Ca^{2+}$ oscillation patterns were induced by using a medium containing 10 mM $Cl^{-}$. The time-laps video represents the same pollen tube as shown in Figure 3B. The images were taken every 5 s for 595 s, assembled as a stack and converted into a video (10 frames/s, 1 s in the video equals to 50 s in the recording).

**Supplemental Table S1: Sequences of oligonucleotide primers**

| Construct | Sequence |
|---|---|
| **Site-directed mutagenesis primers** | |
| CPK21D204A-F | GGTGTGGTTCATCGAGCTCTCAAGCCTGAG |
| CPK21D204A-R | CTCAGGCTTGAGAGCTCGATGAACCACACC |
| CPK23D193A-F | GTGTGATTCATCGAGCTCTCAAGCCTGAG |
| CPK23D139A-R | CTCAGGCTTGAGAGCTCGATGAATCACAC |
| CPK21I373S-F* | GCTCTAAAGGTTAGCGCGGAGAGTCTATC |
| CPK21I373S-R* | GATAGACTCTCCGCGCTAACCTTTAGAGC |
| CPK21I373D-F | GCTAGCTCTAAAGGTTGACGCGGAGAGTCTATC |
| CPK21I373D-R | GATAGACTCTCCGCGTCAACCTTTAGAGCTAGC |
| CPK23S362I-F | GCCCTAAAGGTTATCGCGGTGAGTCTATC |
| CPK23S362I-R | GATAGACTCACCGCGATAACCTTTAGGGC |
| CPK23S362D-F | GCCCTAAAGGTTGACGCGGTGAGTCTATC |
| CPK23S362D-R | GATAGACTCACCGCGTCAACCTTTAGGGC |
| CPK21G2A-F | GATATCGAATTCATGGCTTGCTTCAGCAGTAAACAC |
| CPK21G2A-R | GTGTTTACTGCTGAAGCAAGCCATGAATTCGATATC |
| CPK21G2AC3V-F | GATATCGAATTCATGGCTGTCTTCAGCAGTAAACAC |
| CPK21G2AC3V-R | GTGTTTACTGCTGAAGACAGCCATGAATTCGATATC |
| CPK23G2A-F | GATATCGAATTCATGGCTTGTTTCAGCAGTAAACAC |
| CPK23G2A-R | GTGTTTACTGCTGAAACAAGCCATGAATTCGATATC |
| CPK23G2AC3V-F | GATATCGAATTCATGGCTGTTTTCAGCAGTAAACAC |
| CPK23G2AC3V-R | GTGTTTACTGCTGAAAACAGCCATGAATTCGATATC |
| **Cloning of CDPK-FRET constructs** | |
| CPK21-VK XbaI EcoRI-F | AT<u>TCTAGA</u> <u>GAATTC</u>ATGGGTTGCTTC |
| CPK23-VK XbaI EcoRI-F | AT<u>TCTAGA</u> <u>GAATTC</u>ATGGGTTGTTTC |
| CPK21/23-VK XbaI-R | AT<u>TCTAGA</u>TTCTCCCCCTTTGATCCAAG |
| CPK21-PSCLD ApaI-F | AT<u>GGGCCC</u>GCACCAGACAAGCCTATTG |

| | |
|---|---|
| CPK21-PSCLD SpeI-F | ATACTAGTGCACCAGACAAGCCTATTG |
| CPK21-PSCLD BamHI-F | GGATCCGCACCAGACAAGCCTATTG |
| CPK21-PSCLD SmaI-R | CCCGGGATGGAATGGAAGCAGTTTC |
| CPK21-PSCLD KpnI-R | ATGGTACCATGGAATGGAAGCAGTTTC |
| CPK21-PSCLD SalI-R | ATGTCGACATGGAATGGAAGCAGTTTC |
| CPK21-PSCLD-EF4 SmaI-R | ATCCCGGGCTGCGTGCTGCCACTTC |
| CPK23-PSCLD-EF4 SmaI-R | ATCCCGGGTTGTGTGGTGCCACATC |
| CFP NdeI-F | ATCATATGGTGAGCAAGGGCGAGG |
| CFP ApaI-R | ATGGGCCCCTTGTACAGCTCGTCCATG |
| CPK21 XhoI-F | ATCTCGAGATGGGTTGCTTCAGCAG |
| StrepII SpeI-R | AACTAGTTCATTTTTCGAACTGCGGGTG |

* Used for mutagenesis of I362 in CPK23S362I-CLD21 yielding in CPK23CLD21
F: forward, R: reverse
